# Three cytochrome P450 enzymes catalyse the formation of furanoclerodane precursors in Salvia spp

**DOI:** 10.1101/2024.10.13.618057

**Authors:** Ruoxi Lin, Haixiu Li, Yiren Xiao, Zhuo Wang, Licheng Liu, Gerhard Saalbach, Carlo Martins, Cathie Martin, Evangelos C. Tatsis

## Abstract

Salvia species native to the Americas are rich in valuable bioactive furanoclerodanes, like the psychoactive salvinorin A found in *Salvia divinorum*, which is used in treatment of opioid addiction. However, there is relatively little known about their biosynthesis. To address this, we investigated the biosynthesis of salviarin, the simplest furanoclerodane structure in the ornamental sage *Salvia splendens*. Using a self-organizing map and mutual rank analysis of RNA-seq co-expression data, we identified three cytochrome P450 enzymes responsible for converting kolavenol into salviarin precursors, consecutively: annonene, hardwickiic acid and hautriwaic acid. As annonene and hardwickiic acid have also been proposed as intermediates in the biosynthesis of salvinorin A, and to examine our hypothesis for common evolutionary origin of the furanoclerodane pathway between the two Salvia species, we searched for homologous genes in available data for *S. divinorum*. The enzymes encoded by orthologous genes from *S. divinorum* displayed kolavenol synthase (SdKLS), annonene synthase (SdANS), and hardwickiic acid synthase (SdHDAS) activity respectively, supporting the view that these are intermediate steps in the biosynthesis of salvinorin A. We investigated the origin of annonene synthase and the role of gene duplication in the evolution of this specific activity. Our work shows how *S. splendens* can serve as a model species for studying furanoclerodanes biosynthesis in Salvia species, contributes to understanding the evolution of specialized metabolism in plants, and provides new tools to produce salvinorin A in biotechnological chassis.

## Introduction

Salvia is the largest and the most diverse genus in the family Lamiaceae, with three distinct biogeographical phylogenetic clades in East Asia, the Mediterranean basin and Central Asia, and the Americas (Central and South America) (Walker et al., 2004). Many Salvia species, commonly referred to as sages, are well known for their medicinal properties as well as their culinary and cosmetics applications. The high value of sages has been attributed to their production of bioactive phytochemicals including phenylpropanoids (Exarchou et al., 2002) and terpenoids (Ulubelen, 2003). Several diterpenoids produced by Salvia species have commercial value. For example, the antioxidant food additives carnosic acid and carnosol are abietane diterpenoids produced by culinary European Salvia species (Božić et al., 2015; Ignea et al., 2016; Scheler et al., 2016), the fragrance ingredient sclareol is a labdane diterpenoid produced by *Salvia sclarea* (Schalk et al., 2012) native to the Mediterranean and the tanshinones with cardiovascular-protective properties are a group of abietane diterpenoids, extracted from the roots of the Chinese herb *Salvia miltiorrhiza* (Ma et al., 2021; Wang and Peters, 2022). Half of all Salvia species belong to the subgenus Calosphace, endemic in the neotropics (Central and South Americas) (Ortiz-Mendoza et al., 2022). The potent bioactive substances in this subgenus are furanoclerodane diterpenoids (Li et al., 2016; Ortiz-Mendoza *et al*., 2022; Wu et al., 2012). Clerodanes are a class of diterpenoids with a rearranged labdane carbon skeleton involving the 1,2-shift of two methyl groups. In Lamiaceae, the clerodane scaffold is formed by the paired enzymatic transformation of geranylgeranyl diphosphate (GGPP) to kolavenol, catalysed by class II and class I clerodane synthases in plastids (Li et al., 2023). Following export from the plastid, kolavenol is subjected to several oxidation steps generally catalysed by cytochrome P450 enzymes, with one example reported, so far, in the family Lamiaceae (Kwon et al., 2022).

Clerodanes from the neotropical sages share many structural features such as a furan ring between carbon atoms C13-C16 (Figure 1 - highlighted in blue) and a carboxyl moiety at C18 (Figure 1 - highlighted in orange). Examples of these furanoclerodanes are: salviarins (Fontana et al., 2006b) and salvisplendins (Fontana et al., 2006a) from the ornamental, *Salvia splendens* (scarlet sage) (Dong et al., 2018; Jia et al., 2021); hispanins (Fan et al., 2019) and salvihispins (Fan et al., 2020) from the highly nutritional pseudocereal, *Salvia hispanica* (chia) (Fan *et al*., 2019; Wang et al., 2022a); salvinorins and divinatorins from the Mexican magic sage, *Salvia divinorum* (Siebert, 1994) (Fig. S1). These furanoclerodanes differ mainly in their oxidation sites. The best-known furanoclerodanes are the hallucinogenic salvinorin A and its derivatives from *Salvia divinorum,* which are used in indigenous American folk medicine for pain relief, due to their psychoactive properties (Ortiz-Mendoza *et al*., 2022; Tlacomulco-Flores et al., 2020). Salvinorin A is rather unique among psychedelic substances as it has been identified as a therapeutic for opioid addiction and drug abuse (Kivell et al., 2014). However, understanding of the biosynthesis of these furanoclerodanes is limited, and in places contradictory (Chen et al., 2017; Kwon *et al*., 2022; Ngo, 2019; Pelot et al., 2017).

**Figure 1.**
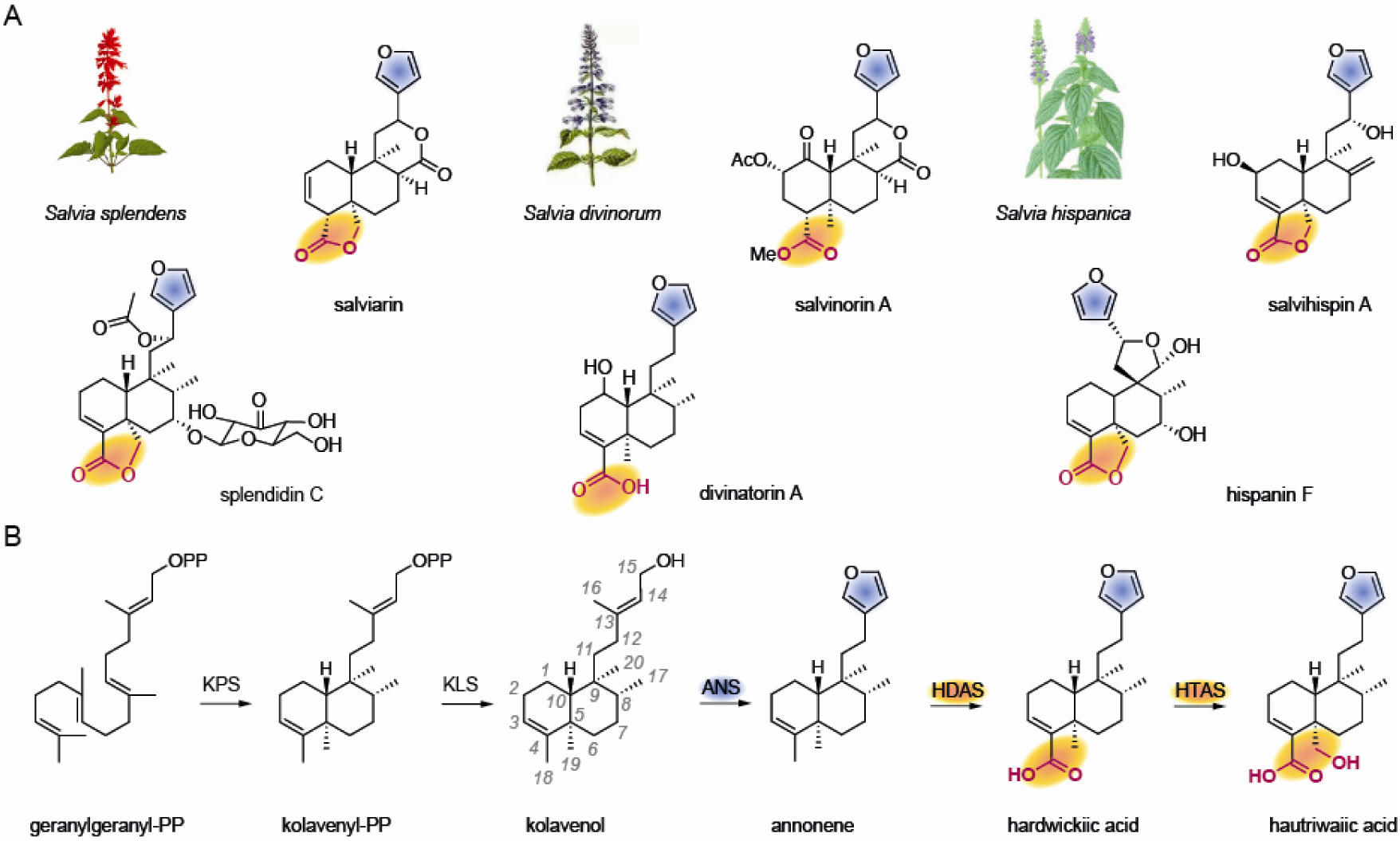
Clerodane metabolism in Salvia species. **A.** examples of furanoclerodanes bearing the furane ring (highlighted in blue) and the oxidized C18, C19 atoms (highlighted in orange) from neotropical sages: the ornamental scarlet sage (*Salvia splendens*), the psychoactive magic sage (*Salvia divinorum*), and the pseudocereal chia (*Salvia hispanica*). **B.** The postulated biosynthesis of furanoclerodanes in Salvia spp (Chen *et al*., 2017). From geranylgeranyl diphosphate (GGPP) to hardwickiic acid (precursor of salvinorin A biosynthesis in *Salvia divinorum*) and to salviarins biosynthesis in *Salvia splendens* through hautriwaic acid as an intermediate. The depicted enzymatic transformations are catalysed by class II diterpene synthase kolavenyl diphosphate synthase (KPS)(Chen *et al*., 2017; Li *et al*., 2023; Pelot *et al*., 2017), class I diterpene synthase kolavenol synthase (KLS)(Li *et al*., 2023), and cytochrome P450 enzymes annonene synthase (ANS), hardwickiic acid synthase (HDAS), and hautriwaic acid synthase (HTAS) (this study).

Here we report the discovery and characterization of genes encoding three cytochrome P450 enzymes acting on the biosynthetic pathway of salviarin in *S. splendens* (Figure 1), based on a combinatorial gene discovery strategy. The combination of two independent gene expression profiling methods with distinct emphasis on gene co-expression patterns streamlined discovery of genes encoding enzymes involved in specialized metabolism. One intermediate, hardwickiic acid, has been identified and isolated from *S. divinorum* and is believed to be involved in salvinorin A biosynthesis (Bigham et al., 2003). To examine our hypothesis that orthologous genes to those of *S. splendens* encoding CYP450 enzymes act in the biosynthesis of salvinorin A in *S. divinorum*, we used sequence homology to search transcriptomic data of *S. divinorum* (Chen *et al*., 2017; Kwon *et al*., 2022; Pelot *et al*., 2017) and identified enzymes with the same catalytic activity. The biochemical and phylogenetic data indicate that in *S. splendens,* the evolution of the clerodane pathway involved recruitment of genes encoding enzymes in the synthesis of other terpenoids. This is evidenced by the neofunctionalization of genes encoding cytochrome P450s with ferruginol synthase activity in other Lamiaceae species to genes encoding annonene synthase operating within the furanoclerodane pathway in both *S. splendens* and *S. divinorum*. This finding aligns with our earlier work on the evolution of genes encoding class II and class I diterpene synthases in the clerodane pathway in *S. splendens* (Li *et al*., 2023). We show that comparison of closely related species with common biosynthetic pathways can contribute to understanding the production of rare, high value fine phytochemicals in plants and opens up new options for their bioproduction.

## Results

### Gene discovery in S. splendens

In earlier work, we identified genes encoding class II and class I diterpene synthases that synthesise kolavenol in *S. splendens* (Li *et al*., 2023). To identify genes encoding the subsequent enzymatic steps in furanoclerodane metabolism in *Salvia splendens*, gene discovery was carried out based on the principle that genes associated with the same biological processes, such as those encoding enzymes in the same metabolic pathway, are often co-expressed. We used the clerodane synthase genes from *S. splendens* as query sequences to select candidate genes from the available *S. splendens* transcriptomic data (Dong *et al*., 2018) (NCBI Accessions: SRX3476236 - SRX3476269). This dataset consisted of three biological replicate samples of tissues including roots, stems, leaves, flower calyxes and corollas from two varieties with red- and purple-coloured flowers; in total, involving thirty different samples. As the first step in transcriptomic analysis, we employed a self-organizing map (SOM) to cluster genes exhibiting similar expression patterns across the different samples into nodes (Figure 2A) using an unsupervised neural network (Dang et al., 2017; Wang et al., 2022b). The formation of clerodane scaffolds is catalysed by the class II diterpene synthase with kolavenyl diphosphate synthase (KPS) activity encoded by the gene *SspdiTP2.1* (Li *et al*., 2023). In *S. splendens* the conversion of kolavenyl diphosphate to kolavenol is the second step in clerodane biosynthesis, catalysed by class I diterpene synthases (diTPSs). These class I diTPSs are either specific for kolavenol synthesis (SspdiTPS1.5), or they also possess kaurene synthase activity (KS; genes *SspdiTPS1.1 & SspdiTPS1.2*) or miltiradiene synthase activity (MS; gene *SspdiTPS1.3*)*, in vitro* (Li *et al*., 2023).

**Figure 2.**
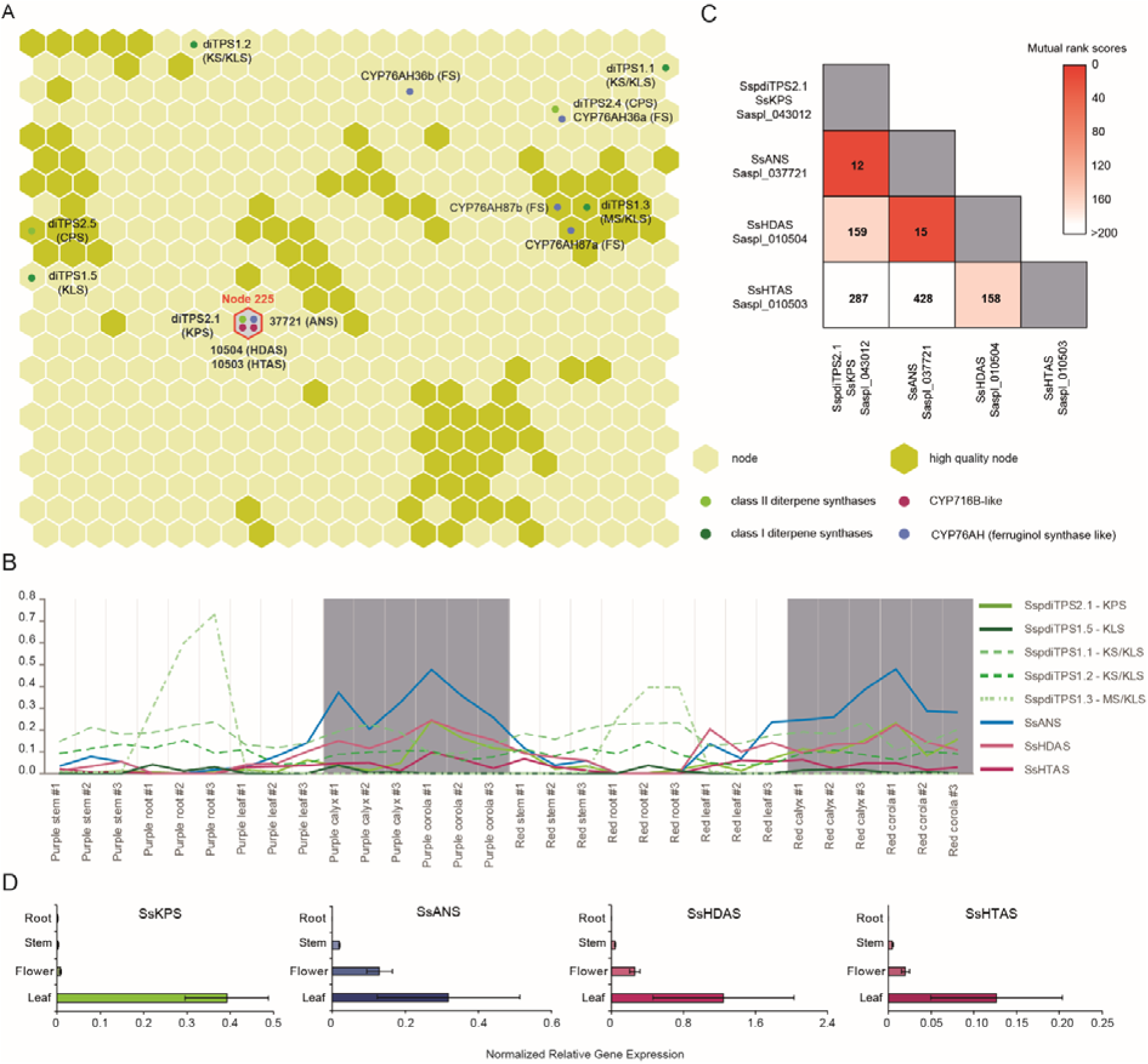
Transcriptomic data analysis of *Salvia splendens*. **A**, Self-organizing map of *S. splendens* transcriptomic data. Each hexagon node represents at least 70 genes with the most similar expression profiles. Darker coloured nodes represent the higher quality nodes. The placement in nodes of previously identified diterpene synthase genes (Li *et al*., 2023) are highlighted in light (class II diTPS) and dark (class I diTPS) green dots and cytochrome P450 enzymes of CYP76AH subfamily in blue dots. The enzymatic activity is shown in brackets. The class II clerodane synthase gene *SsKPS* is placed in node 225 (highlighted by an orange perimeter) together with annonene synthase gene *SsANS* (CYP76AH-*like*), hardwickiic acid synthase gene *SsHDAS* (CYP716B-like) and hautriwaic acid synthase gene *SsHTAS* (CYP716B-like). **B**, Expression profiles of genes encoding enzymes with SsKPS, SsKLS, SsANS, SsHDAS and SsHTAS activity across 30 tissues. **C,** Heat map of the co-expression analysis of *SsKPS*, *SsANS*, *SsHDAS* and *SsHTAS* in *S. splendens* transcriptomic data. A low MR ranking indicates a relatively similar expression profile. **D**, qRT-PCR analysis of the expression for *SsKPS*, *SsANS*, *SsHDAS* and *SsHTAS* genes from different tissues (root, stem, flower, leaf) of *S. splendens*. Data were obtained from three independent biological replicates. Transcript levels were normalized to that of actin (n =3). Bars represent SD.

As a starting point, we used the kolavenyl diphosphate synthase (KPS) gene (*SspdiTPS2.1/SsKPS Saspl_043012*) (Li *et al*., 2023) as our “bait”. This gene encodes the first step in the formation of the clerodane skeleton, which is part of node 225 (Figure 2A) comprising a total of 78 genes. The average expression level for genes in node 225 was higher in leaves and reproductive tissues (flower corolla and calyx) and lower in roots (Figure 2B, Fig. S2). Among these genes twelve were functionally annotated as encoding cytochrome P450 enzymes. To prioritize the choice of cytochrome P450 genes for cloning, expression and activity assays, we additionally applied mutual rank-based co-expression analysis. Mutual ranking analysis (MRA) of the transcriptome data involved the pairwise comparison of gene expression levels across different samples (Obayashi and Kinoshita, 2009). MRA ranked input genes based on the similarities in their co-expression patterns against the query genes, and provided rankings (Ding et al., 2019).

### Discovery and characterization of Annonene Synthase (SsANS)

To screen for candidate P450 genes involved in clerodane biosynthesis, we assembled the kolavenol biosynthetic pathway (Figure 3A) (Li *et al*., 2023) in baker’s yeast (*Saccharomyces cerevisiae*). The geranylgeranyl diphosphate synthase (*ERG20*::F96C) (Ignea et al., 2014) and a fused class II (*SsKPS*; *SspdiTPS2.1*) and class I (*SsKLS*; *SspdiTPS1.5*) clerodane synthase gene (Li *et al*., 2023; Zhou et al., 2012) were cloned into pESC-His vector for recombinant expression in yeast. Using both SOM and MRA rankings, we focused on cytochrome P450 genes that are part of the SOM node 225 and are highly ranked by MRA in relation *to SsKPS*. Our top candidate gene *Saspl_037721* was placed in node 225 of SOM and was ranked as 12^th^ most highly co-expressed gene with *SsKPS* (*Saspl_043012*) among all genes in MRA (Figure 2C). *Saspl_037721* is located on genomic scaffold 129 (Dong *et al*., 2018) and is functionally annotated as a ferruginol synthase-like (CYP76AH) gene. We cloned the full-length transcript of *Saspl_037721* into the pESC-Leu vector together with a cytochrome P450 reductase gene (*CrCPR2*) from *Catharanthus roseus* (Wang *et al*., 2022b; Wang et al., 2022c) and transformed into yeast strain AM119 with the engineered pESC-His vector. After galactose induction, the recombinant yeast culture was extracted with ethyl acetate and analysed by GC-MS. A new peak appeared at 33.8 min (Figure 3B, Fig. S3A) under the extracted ion chromatogram (EIC) mode at *m/z* 271. Using the mass spectrum of the new peak to search the NIST17 database, annonene was the best match (75.2% probability) (Figure 3C, Fig. S3B-E). Since Saspl_037721 encoded a protein with anonnene synthase activity, hereafter it will be referred to as annonene synthase (SsANS).

**Figure 3.**
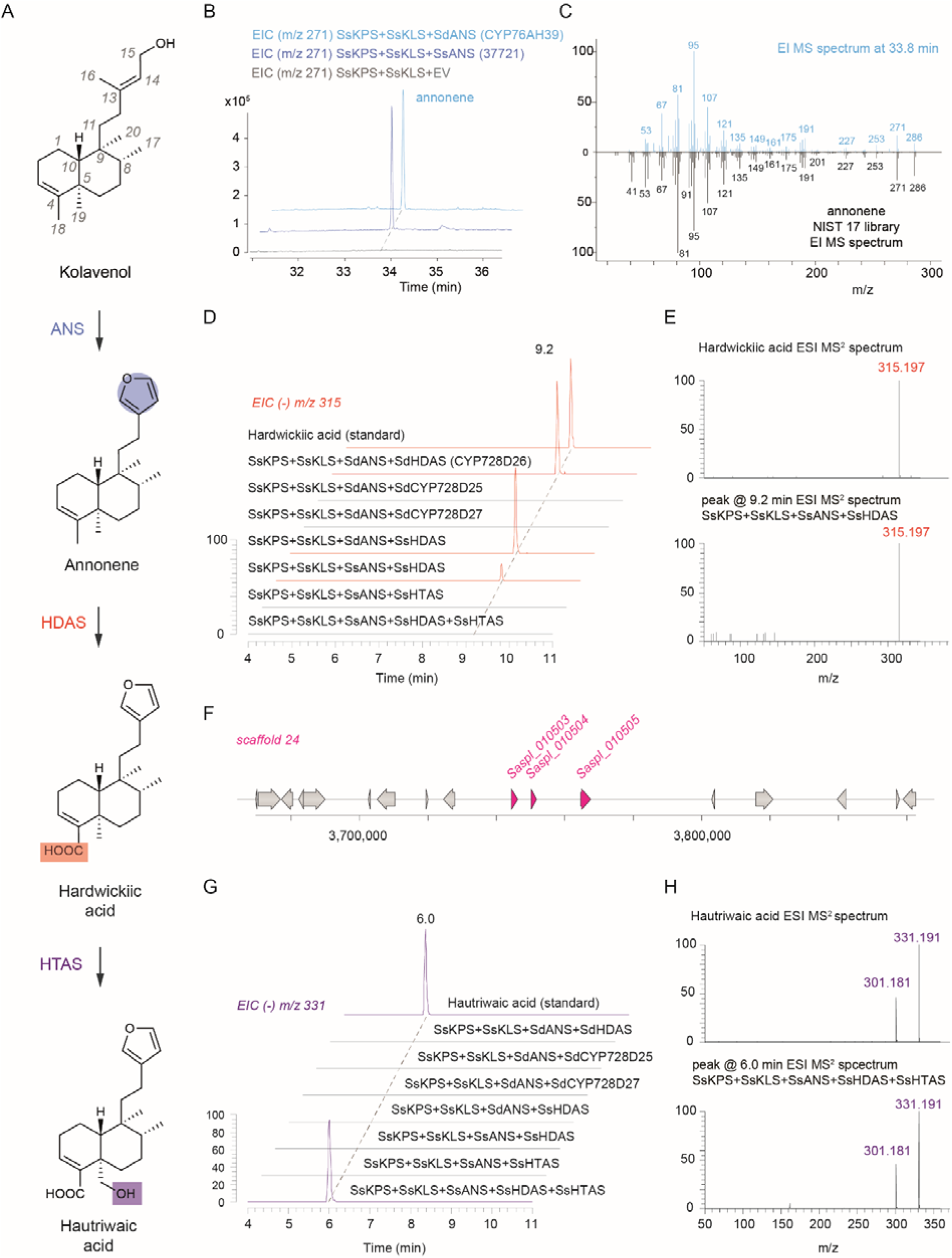
Characterization of enzymes in the clerodane diterpenoid biosynthetic pathway from kolavenol to hautriwaic acid. **A**, The biosynthetic pathway from kolavenol to hautriwaic acid. **B**, GC-MS analysis (selected *m/z* signal of 271) of yeast strains expressing GGPPS (Erg20::F96C), SsKPS, SsKLS, SsANS or SdANS, and CrCPR in blue. Negative control (in grey): yeast strain expressing GGPPS (Erg20::F96C), SsKPS, SsKLS and CrCPR. **C**, Comparison of EI mass spectrum of GC-MS peak eluting at 33.8 min (panel B – product of SdANS) with the annonene standard (EI mass spectrum from NIST 17 library). **D**, LC-MS analysis (selected *m/z* signal of 315 in negative mode) of yeast strains expressing GGPPS (Erg20::F96C), SsKPS, SsKLS, SsANS, CrCPR and combinations of SsHDAS or SdHDAS, SsHTAS. **E**, Comparison of ESI mass spectrum of LC-MS peak eluting at 9.6 min (panel D) with the hardwickiic acid standard. **F**, Genomic area (scaffold 24) from *S. splendens* genome showing that *SsHDAS* (*Saspl_010504*) and *SsHTAS* (*Saspl_010503*) genes are the result of tandem duplications. **G**, LC-MS analysis (selected *m/z* signal of 331 at negative mode) of yeast strains expressing GGPPS (Erg20::F96C), SsKPS, SsKLS, SsANS or SdANS, CrCPR and combinations of SsHDAS or SdHDAS, SsHTAS. **H**, Comparison of ESI mass spectrum of LC-MS peak eluting at 6.7 min (panel G) with the hautriwaic acid standard.

### Discovery and characterization of Hardwickiic Acid Synthase (SsHDAS) and Hautriwaic Acid Synthase (SsHTAS)

We performed MRA with reference to *SsANS* to select cytochrome P450 gene candidates that might encode proteins acting on annonene. The *Saspl_010504* gene was annotated as encoding a cytochrome P450 716B1-like protein, ranked 15^th^ by MRA in relation to *SsANS* and clustered in node 225 in SOM like Ss*KPS* and *SsANS*. We cloned *Saspl_010504* into the pESC-Ura vector and tested its activity *in vivo* following galactose induction in recombinant yeast. The ethyl acetate extract of yeast was analysed by GC-MS, and a new peak at EIC *m/z* 250 was detected eluting at 36.9 min (Fig. S4A-B). Searching the mass spectra in the NIST17 database suggested that the new peak had a top match with hardwickiic acid (79.3% probability, Fig. S4C). Since hardwickiic acid bears a carboxylic acid moiety, we shifted our analysis to LC-MS. The hardwickiic acid standard eluted at 9.2 min with *m/z* = 315 in negative mode, which aligned with the peak of the Saspl_010504 enzyme product in the assay (Figure 3D) and agreed with the MS^2^ spectrum (Figure 3E, Fig. S5). Therefore, we concluded that the cytochrome P450 encoded by *Saspl_010504* acts on anonnene and functions as a hardwickiic acid synthase (SsHDAS).

The gene *Saspl_010503* is a tandem duplicate of *Saspl_010504* (Figure 3F), the encoded proteins sharing high sequence similarity at the amino acid level (83.4%) and it is also functionally annotated as a CYP716B1-like cytochrome P450. *Saspl_010503* was also placed in node 225 of SOM with a similar expression profile to *SspdiTPS2.1 (SsKPS)*, *SsANS* and *SsHDAS* (Figure 2A). Despite the low-ranking in MRA (Figure 2C), we tested whether *Saspl_010503* encodes a protein acting in the pathway. When we co-expressed the cytochrome P450 encoded by *Saspl_010503* together with SsKPS, SsKLS and SsANS, no hardwickiic acid or any other product was formed with annonene by LC-MS analysis (Figure 3D, Fig. S5). When we co-expressed the cytochrome P450 encoded by *Saspl_010503* with SsKPS, SsKLS, SsANS and SsHDAS, a new peak appeared in LC-MS at 6.7 min with an *m/z* 331 (negative mode) (Figure 3G, Fig. S6). The new product was identified as hautriwaic acid based on comparison of its retention time and MS^2^ spectrum with an authentic standard (Figure 3G/H, Fig. S6), and therefore Saspl_010503 was named as hautriwaic acid synthase (SsHTAS).

### Expression of genes encoding cytochrome P450s acting in clerodane biosynthesis

We conducted qRT-PCR to assess the expression level of all genes associated with hautriwaic acid biosynthesis in SOM node 225. The measurements covered various plant tissues, including roots, stems, leaves, and flowers (Figure 2D). These clerodane biosynthesis genes exhibited significantly elevated expression levels in flowers and leaves, which aligned with metabolomic data showing the predominant accumulation of *S. splendens* clerodanes, such as salviarin, in these tissues (Fig. S7).

### Key clerodane pathway intermediates accumulate in abaxial peltate trichomes

In *S. divinorum,* salvinorin A and other furanoclerodanes are specifically accumulated in peltate trichomes on the abaxial side of the leaves (Chen *et al*., 2017; Siebert, 2004b). *S. splendens* leaves have both capitate and peltate glandular trichomes (Agustin et al., 2022; Ghonam et al., 2014; Perveen). SEM confirmed that *S. splendens* leaves predominantly have peltate glandular trichomes up to 50 μm in diameter on the abaxial side (Figures 4, Fig.S8A). Metabolite analysis of these trichomes, picked from the abaxial side of the leaves of *S. splendens*, showed the presence of annonene, hardwickiic acid, hautriwaic acid and salviarin (Figure 4, Fig. S8B). The results suggested that the furanoclerodanes in *S. splendens* are produced within glandular trichomes, similar to the production of salvinorin A in *S. divinorum*. Additionally, we undertook matrix-assisted laser desorption/ionization (MALDI) mass spectrometry imaging (MSI) on the abaxial side of *S. splendens* leaves (Figures 4A-B) to identify the site of salviarin accumulation (Figure 4). MALDI-MSI displayed the co-presence of annonene (Figure 4C), hardwickiic acid (Figure 4D), and hautriwaic acid (Figure 4E) with salviarin (Figure 4F) at the same positions with the peltate trichomes (Figure 4G).

**Figure 4.**
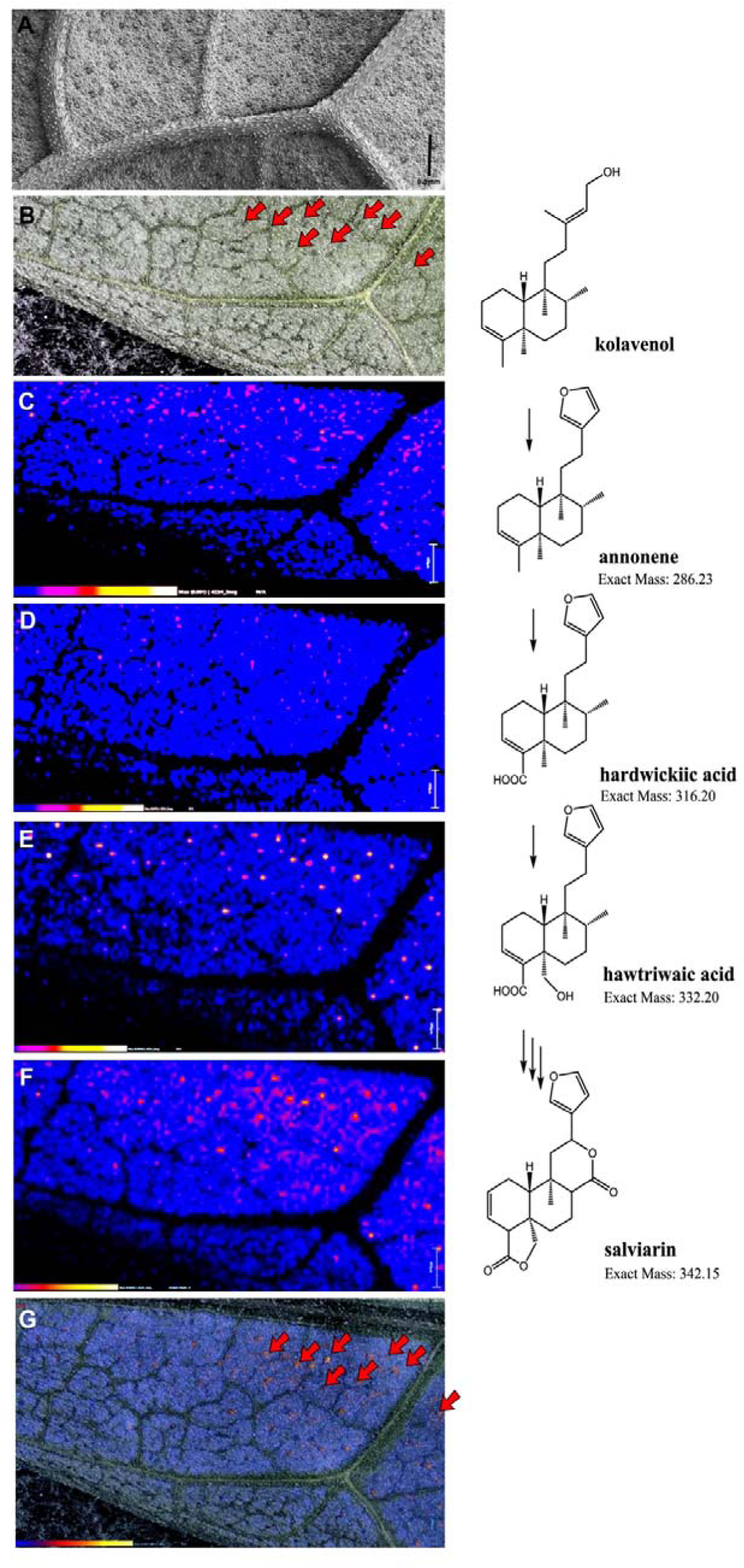
Identification of intermediates in the pathway synthesizing salviarin on the abaxial surface of a leaf of *Salvia splendens* by MALDI-imaging. **A**, Scanning electron micrograph (SEM) of abaxial surface of leaf of *S. splendens*. A magnification of part of this image is shown in supplementary Fig. S8A. This is not the same leaf that was used for MALDI-imaging in panels C-G. **B,** Photograph of abaxial surface of a leaf of *S. splendens*. The red arrows indicate some of the sites of medium-sized peltate trichomes (shown in detail in the SEM in Supplementary figure 8A). **C,** MALDI image of annonene (mass 285 in negative ion mode; scale 0-0.001, scale bar = 720 μm) **D,** MALDI image of hardwikiic acid (mass 315 in negative ion mode, scale 0-0.001, scale bar = 452 μm). **E,** MALDI image of hawtriwaic acid (mass 331 in negative ion mode, scale 0-0.0034, scale bar = 452 μm). **F,** MALDI image of salviarin (mass 341 in negative ion mode, scale 0-0/0045, scale bar = 452μm). Note the blurring of the salviarin signal suggesting it may have been released from the trichomes and spread over the leaf surface. **G,** Overlay of panel F on panel B to show the coincidence of the salviarin signal with the peltate trichomes on the abaxial leaf surface. The red arrows highlight the same trichomes shown in panel B. The intensity of the signal for each compound is shown by the scale bars at the bottom of each image.

### Identification and characterization of kolavenol synthase, annonene synthase and hardwickiic acid synthase in Salvia divinorum

Convergence is common in plant specialised metabolism and despite many phytochemicals having identical or very similar chemical structures, it cannot be assumed that they are synthesised the same way, biologically. In work by Chen *et al*. (2017), a scheme for the biosynthesis of salvinorin A and other clerodanes was proposed with annonene and hardwickiic acid as intermediates between kolavenyl diphosphate (the product of the class II clerodane synthase) and salvinorin A. Detection and quantification of hardwickiic acid by LC-MS in peltate trichomes from *S. divinorum* (Siebert, 2004a) also supported its role as potential intermediate in furanoclerodane metabolism. Considering that both *S. splendens* and *S. divinorum* belong to the Calosphace subgenus and originate from the same geographical region producing highly similar furanoclerodanes, we tested whether the early steps in furanoclerodane biosynthetic pathways share functionally equivalent enzymes between these two species. In previous work on *S. divinorum,* the class II clerodane synthase, kolavenyl diphosphate synthase (KPS), was characterized based on its expression in peltate trichomes and transcriptomic data from peltate trichomes on the abaxial leaf surface (Chen *et al*., 2017; Pelot *et al*., 2017). However, attempts to identify a functional class I diTPS with kolavenol synthase activity have not yet been successful (Chen *et al*., 2017; Pelot *et al*., 2017).

To examine the relevance of genes encoding cytochrome P450 enzymes in furanoclerodane metabolism from *S. splendens* to *S. divinorum*, we searched transcriptomic data from *S. divinorum* peltate trichomes (Pelot *et al*., 2017) (NCBI accession number SRX1875790) for candidate genes encoding proteins involved in salvinorin A biosynthesis by sequence similarity. We searched first for genes encoding class I diterpene synthases closely related to those characterized functionally with kolavenol synthase activity in *S. splendens*; SspdiTPS1.5 which acts specifically as a kolavenol synthase, SspdiTPS1.1 & SspdiTPS1.2 which also have kaurene synthase activity, and SspdiTPS1.3 which also has miltiradiene synthase activity. Three transcripts encoding three previously reported class I diTPSs came top in the results. The first enzyme, called SdKSL1, aligned most closely to SspdiTPS1.3 (Fig. S9); a second enzyme, called SdKSL2 was most closely aligned to SspdiTPS1.5 (Fig. S9), while a third one called SdKSL3 (Pelot *et al*., 2017) aligned closely with SspdiTPS1.1 and SspdiTPS1.2 (Fig. S9). Both SdKSL2 and SdKSL3 were highly expressed in peltate trichomes of *S. divinorum* (Chen *et al*., 2017; Pelot *et al*., 2017). Pelot *et al* reported that neither of these class I diTPS genes had kolavenol synthase activity in *in vivo* assays dependent on a class II KPS activity (SdCPS2). Given the limitations of *in vivo* assays due to the activity of endogenous phosphatases in *N. bentamiana*, it seemed likely that, by analogy to *S. splendens*, SdKSL2 and SdKSL3 might provide the class I diterpene synthase activity for salvinorin A biosynthesis.

The genes *SdKSL1* and *SdKSL2* were synthesized to encode truncated proteins lacking their transit peptides and were expressed in *E. coli* (Li *et al*., 2023). To characterize the activity towards kolavenyl diphosphate of SdKSL1 and SdKSL2, a coupled *in vitro* enzyme assay was performed with GGPP and SsKPS (Li *et al*., 2023). Both SdKSL1 and SdKSL2 were able to convert the kolavenyl diphosphate (product of SsKPS) to kolavenol and could potentially act as kolavenol synthases in clerodane metabolic pathway in *S. divinorum* (Figure 5A and B). Our searches of the *S. divinorum* transcriptome from peltate trichomes (Pelot *et al*., 2017) identified a transcript, Sd_DN33_c0_g1_i12 encoding a protein most closely related to SsANS (Fig. S11). Sd_DN33_c0_g1_i12 is identical to the *SdCS* gene which encodes the cytochrome P450 CYP76AH39 with reported crotonolide G synthase activity, while it shares 82.3% amino acid sequence similarity to *S. splendens* SsANS (Kwon *et al*., 2022). In *S. splendens*, we could not detect the presence of crotonolide G in the peltate trichomes on the abaxial side of the leaf. We synthesised the Sd_DN33_c0_g1_i12 transcript, expressed it and tested the enzymatic activity together with SsKPS and SsKLS in yeast. GC-MS analysis of the *in vivo* assay showed a chromatographic peak in EIC *m/z* 271 eluting at 33.8 min (Figure 3B), the same as the annonene produced by SsANS, while a search of the mass spectrum in the NIST17 database returned a score of 90.3% probability of annonene (Figure 3C). To prove the annonene synthase activity of Sd_DN33_c0_g1_i12, we paired Sd_DN33_c0_g1_i12 with SsHDAS in yeast. LC-MS analysis of the products of the *in vivo* assay showed the elution of a peak at 9.2 min with MS and MS^2^ identical to the hardwickiic acid standard (Figure 3D, Fig. S5), and therefore we refer to Sd_DN33_c0_g1_i12 as SdANS.

**Figure 5.**
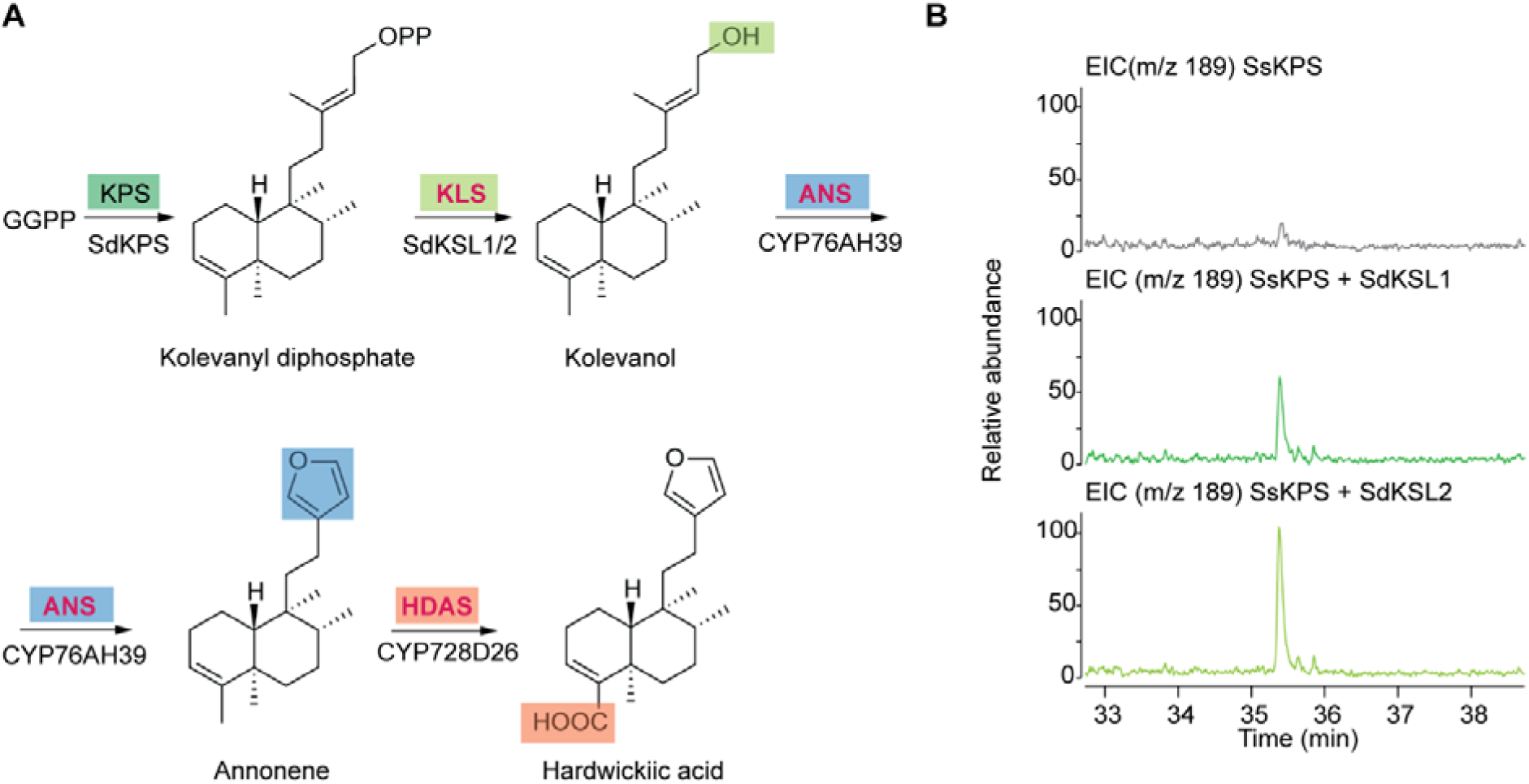
Characterization of enzymes with kolavenol synthase (KLS) and the furanclerodane biosynthesis in *Salvia divinorum.* **A,** The enzymatic steps in biosynthesis of hardwickic acid in *S. divinorum* starting from geranyl-geranyl diphosphate (GGPP). On a coloured background are depicted the enzymatic activities, with bold red fonts marking the enzymatic activities characterized in the present study (KPS: kolavenyl diphosphate synthase performed by SdKPS1 (Chen *et al*., 2017; Pelot *et al*., 2017); KLS: kolavenol synthase performed by SdKSL1-2 (genes reported but not functionally characterised as KLS by (Chen *et al*., 2017; Pelot *et al*., 2017)); ANS: annonene synthase (Figure 3B) performed by CYP76AH39 (gene reported but not functionally characterised as ANS by (Kwon *et al*., 2022)); HDAS: hardwickiic acid synthase (Figure 3D) performed by CYP728D26 (gene reported but not functionally characterised as HDAS by (Kwon *et al*., 2022));. **B**, GC-MS analysis (selected m/z signals at 189) of extracts from enzyme assays against GGPP of class II diterpene synthase SsKPS with combinations of class I diterpene synthases, SdKSL1 and SdKSL2. Both class II and class I diterpene synthase enzymes were heterologously expressed and purified from *E. coli.* The enzyme assay of SsKPS without the addition of class I enzymes or alkaline phosphatase was used as a negative control.

Using SsHDAS as a search template, we identified a 1130 bp partial transcript sequence, Sd_DN6570_c0_g1_i3, in the *S. divinorum* trichome transcriptome, and the full length of this transcript had been deposited by (Kwon *et al*., 2022) under the name CYP728D26 (NCBI accession: QMS79245.1). Another top hit from the search results, Sd_DN990_c0_g1_i18, was a transcript aligned partially to the sequences of *CYP728D25* and *CYP728D27* (NCBI accessions: QMS79244.1 and QMS79246.1), deposited by the same authors (Fig. S12). These three genes were reported to encode candidate enzymes involved in salvinorin A biosynthesis, based on their presence in the S. *divinorum* peltate trichome-specific transcriptomic data (14). CYP728D25, CYP728D26, CYP728D27 have nucleotide sequence similarities of 83.6%, 85.9% and 83.3% to *SsHDAS* respectively and amino acid sequence similarities of 71.5%, 75.6% and 71.5%, respectively to SsHDAS (Fig. S12). We synthesised the three genes and expressed them together with *S. splendens* SsKPS, SsKLS and *S. divinorum* SdANS to assess their *in vivo* enzymatic activity. Our analysis of ethyl acetate extracts of yeast by LC-MS showed that only CYP728D26 exhibited HDAS activity, as evidenced by the appearance of a new peak at EIC *m/z*=331 (Figure 3D, Fig. S5), aligning with the standard of hardwickiic acid. Therefore, we refer to CYP728D26 as SdHDAS.

### The gene encoding annonene synthase evolved from ferruginol synthase

In the latest chromosome level *S. splendens* genome assembly, the tandemly duplicated genes, *Saspl_037721* and *Saspl_037719*, lie on chromosome 19 in proximity to *SspdiTPS1.3* and the non-functional class II diterpene synthase gene *SspdiTPS2.8* (Table S1; Li et al., 2023). The genome of *S. splendens* has been shaped by two relatively recent whole genome duplication events (WGD). The older WGD was shared by *S. hispanica* and *S. splendens,* while the more recent one is unique to *S. splendens* (Wang *et al*., 2022a). Thus, in an intraspecies syntenic analysis, the genomic region bearing *SsANS* has three other syntenic regions on chromosomes 10, 20 and 21 in *S. splendens*. The syntenic relationships between class II diterpene synthases *SspdiTPS2.8* and *SspdiTPS2.4* from chromosome 10, *SspdiTPS2.5* from chromosome 21, and *SspdiTPS2.2* from chromosome 20 are highlighted by green ribbon in Figure 6. Both SspdiTPS2.4 and SspdiTPS2.5 enzymes have been characterised as copalyl diphosphate synthases (Li *et al*., 2023). The dark green ribbon shows the synteny of genes encoding class I diterpene synthases; *SspdiTPS1.3* from chromosome 19 and *SspdiTPS1.4* from chromosome 20. *SsANS* and its tandem duplicate *Saspl_037719* exhibit a syntenic relationship (highlighted by the navy-blue ribbon, Figure 6B) with cytochrome P450 genes *CYP76AH36a* and *CYP76AH36b* encoding ferruginol synthase activity. Additionally, the characterized ferruginol synthase gene CYP76AH87b has a syntenic relationship only to the *CYP76AH87a* gene (highlighted by the pale blue ribbon) (Li *et al*., 2023). We have reported a ferruginol biosynthetic gene cluster (BGC) in the two biggest subfamilies of the family Lamiaceae, Nepetoideae and Scutellaroideae, with *SspdiTPS2.4*, *SspdiTPS2.5*, *SspdiTPS1.3, SspdiTPS1.4*, *CYP76AH36a/b, CYP76AH87a/b* all being part of the ferruginol BGC in *S. splendens* (Li *et al*., 2023). We used an engineered yeast strain producing miltiradiene (Li *et al*., 2023) to assay SsANS and Saspl_037719 against miltiradiene, but no ferruginol synthase activity was observed (Figure 6C). When we assayed CYP76AH36a/b and CYP76AH87a/b against kolavenol, annonene synthase activity was detected for both enzymes (Figure 6A). The phylogeny of syntenous cytochrome P450s from Lamiaceae (Fig. S13A) shows that the gene encoding SsANS diverged from that encoding CYP76H87 around 17.8 MYA, and the divergence of the gene encoding SsANS from the syntenous CYP76AH36 genes with ferruginol synthase activity occurred around 10.6 MYA (Fig. S13B). These enzymatic activities, phylogenetic and syntenic relationships suggest that the emergence of SsANS was the result of neofunctionalization of an ancestral gene encoding f erruginol synthase and this shift in activities is parsimonious with other reported examples of neofunctionalization in plant specialised metabolism (Weng, 2014).

**Figure 6.**
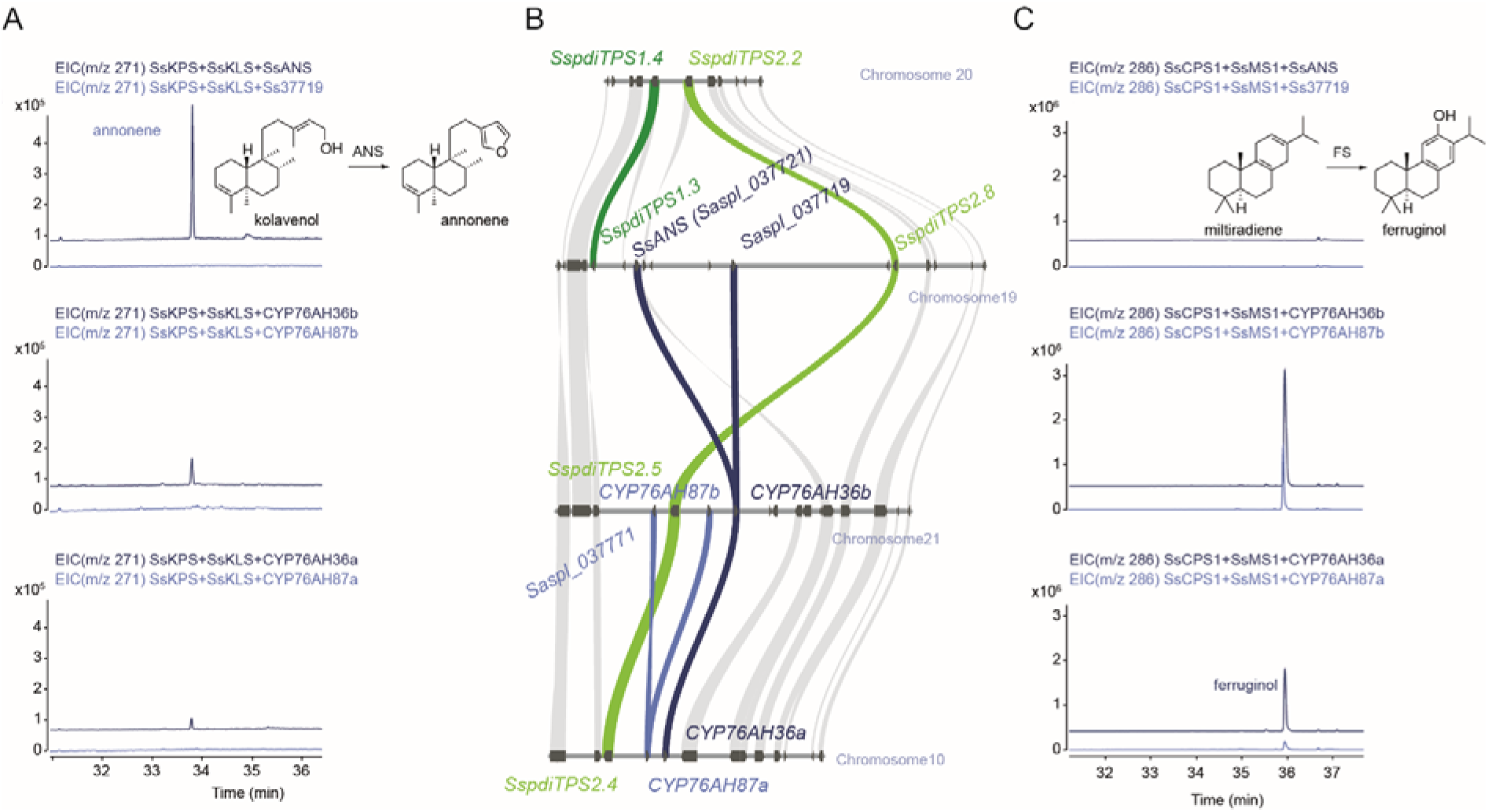
Neofunctionalization of annonene synthase SsANS from ferruginol synthase. **A**, GC-MS analysis (selected ion chromatogram at *m/z* 271) of yeast extracts expressing cytochrome P450 enzymes together with class II (kolavenyl diphosphate synthase – KPS) and class I (kolavenol synthase – KLS) clerodane synthases from *S. splendens*. **B**, Intraspecies syntenic analysis of the regions of the *S. splendens* genome containing genes encoding SsANS, ferruginol synthase, class II and class I diTPSs. The light green ribbon highlights the syntenic relationships among class II diTPSs, dark green ribbon highlights the class I diTPSs, light blue ribbon highlights the CYP76AH87s P450s and the dark blue ribbon highlights the CYP76AH36s P450s. **C**, GC-MS analysis (selected ion chromatogram at *m/z* 286) of yeast extracts expressing cytochrome P450 enzymes together with copalyl diphosphate synthase (CPS) and miltiradiene synthase (MS).

### Biosynthesis of furanoclerodanes in the subgenus Calosphace likely evolved after subgenus divergence

The class II and class I diterpene synthases operating in the clerodane pathway in the family Lamiaceae were recruited from gibberellin and abietane biosynthetic pathways, respectively (Li *et al*., 2023). Phylogenomic analysis supported by biochemical data have demonstrated the polyphyletic origin of clerodane biosynthetic pathways in *Scutellaria barbata* and *S. splendens* through repeated evolution with the latter evolved more recently. The estimated time of divergence of SsKPS from SdKPS is around 20 million years ago (MYA) and for SsKLS about 15 MYA (Li *et al*., 2023). The divergence time of *S. splendens* from its Eurasian relatives is estimated to have been 23.8 MYA (Fig. S14) while the speciation time between *S. splendens* and *S. hispanica* is estimated to have been 6.4 MYA (Fig. S14). The lack of any syntenic relationships between the genes encoding CYP76AH36 cytochrome P450 enzymes and the CYP genes *from S. miltiorhiza* may reflect the absence of orthologues in Eurasian Salvias while the syntenic counterparts in *S. hispanica* may be encoded by genes orthologous to those in *S. splendens* (Fig. S13A).

Phylogenomic analyses indicate that the *SsHDAS* and *SsHTAS* in *S. splendens* have orthologous counterparts in *S. hispanica* (Fig. S12, Fig. S15). Estimation of divergence times based on Bayesian phylogeny suggests the divergence between SsHDAS and SsHTAS occurred around 9.4 MYA (Fig. S16), indicating the emergence of enzymes with HDAS and HTAS activities before the speciation of *S. splendens* and *S. hispanica*. Therefore, the early steps (ANS and HDAS) of furanoclerodane biosynthesis presumably utilized orthologous genes encoding enzymes that catalysed the same enzymatic reaction, with minimal or no alternative routes. Although precise estimates of the speciation time for *S. divinorum* are unavailable due to lack of genome sequence, given that Calosphace is a monophyletic subgenus, all species within this subgenus are closely related (Walker *et al*., 2004). The identification of *SdANS* and *SdHDAS* using homology-based search supports the hypothesis that the emergence of ANS and HDAS are two evolutionary events shared by the neotropical sages of the subgenus Calosphace.

## Discussion

To understand better furanoclerodane biosynthesis in *S. splenden*s, we initially employed an unbiased and unsupervised neural network self-organizing map to cluster all the genes from *S. splendens* into different nodes, based on gene expression profiles. Both class I and class II clerodane synthases were used as guides to focus on specific nodes, although genes encoding proteins with kolavenol synthase activity (class I) were not precisely co-expressed with KPS or the three cytochrome P450 genes acting in the furanoclerodane pathway in different tissues, possibly reflecting the observed catalytic redundancy of kolavenol synthase (Li *et al*., 2023). The bifunctional class I diterpene synthases SspdiTPS1.1 and SspdiTPS1.2 (Li *et al*., 2023) with both kaurene and kolavenol synthase activity were expressed in all plant tissues analysed (Figure 1) and likely fulfil the role of kolavenol synthase, although their expression profiles were not co-ordinate with those of *SsKPS* or *SsANS, SsHDAS* or *SsHTAS*. The clerodane-specific KLS gene (*SspdiTPS1.5*) showed very low expression in all tissues although this could reflect expression in very specialised cell types such as trichomes on leaves. Hypothesizing that cytochrome P450 enzymes oxidize the clerodane skeleton, we employed MRA to prioritize the selection of genes encoding cytochrome P450 enzymes for enzyme assays. This strategy led to the discovery of three cytochrome P450 enzymes involved in furanoclerodane biosynthesis in *S. splendens,* forming annonene, hardwickiic acid and hautriwaic acid which we propose are intermediates in salviarin biosynthesis. (-)-Hardwickiic and hautriwaic acid have been reported as constituents of neotropical *Salvia wagneriana* (Bisio et al., 2004) and desert hybrid *Salvia* x *jamensis* (Bisio et al., 2009). Both (-)-hardwickiic and hautriwaic acid have been suggested as drug design probes for therapies impacting pathologies of the central nervous system (Pittaluga et al., 2013), including relief for neurological pain and visceral hypersensitivity (Cai et al., 2019). (-)-Hardwickiic acid is also used commercially as an antagonist of bitter taste receptors to mask or decrease perception of bitterness and/or to enhance perception of sweetness (WO2013072332A1, 2013).

The best-known example of a furanoclerodane from Savia species is salvinorin A from *S. divinorum* which is known for its use in the treatment of psychiatric diseases (Li *et al*., 2016; Ortiz-Mendoza *et al*., 2022; Wu *et al*., 2012). As furanoclerodanes from *S. splendens* share many of the chemical features of salvinorin A which are essential for its anti-opioid activity, such as the furan ring and the carboxyl group at C18 on the clerodane scaffold (Prisinzano and Rothman, 2008; Roach and Shenvi, 2018), it is likely to share the same furanocleordane biosynthetic enzymes as its close relative *S. divinorum.* Our divergence time analysis performed on syntenic P450s between *S. splendens* and another Calosphace species, *S. hispanica*, indicates that the initial steps of clerodane biosynthesis probably are the same and share a common ancestor within this subgenus. Indeed, hardwickiic acid has been isolated from *S. divinorum* (Bigham *et al*., 2003; Chen *et al*., 2017), and shown to be co-localised in peltate glandular trichomes on the abaxial surface of the leaves of *S. divinorum* (Chen *et al*., 2017), leading to the proposal of a biosynthetic route for salvinorin A with annonene and hardwickiic acid intermediates (Chen *et al*., 2017). However, difficulties in identifying class I diTPS activity in *S. divinorum* led to the suggestion that crotonolide G was the first committed intermediate in furanoclerodane biosynthesis, and the identification of cytochrome P450 enzymes with decorating activity on crotonolide G was reported (Kwon *et al*., 2022; Ngo, 2019). We used the class I diTPSs and cytochrome P450 enzymes from *S. splendens* to search for transcripts encoding orthologues in *S. divinorum* through homology alignment of the encoded proteins. Genes encoding orthologues of the class I diTPSs, SsANS and SsHDAS from *S. splendens* were identified from the transcriptomic data from peltate trichomes producing salvinorin A in *S. divinorum* and these were characterized by *in-vitro* and *in-vivo* enzyme assays. The identified SdKSL1/2, SdANS and SdHDAS enzymes from *S. divinorum* displayed the same activities as their orthologues in *S. splendens*. We used transit peptide cleavage sites (Fig. S10) different to earlier studies to express the proteins (Chen *et al*., 2017; Pelot *et al*., 2017) and identified the activity of SdKSL1 and SdKSL2. This difference may explain why no kolavenol synthase enzymatic activity was detected in earlier studies. Despite the fact that SdANS (CYP76AH39) has been reported to act on clerodane metabolism as a crotonolide G synthase, our *in vivo* assays suggested it produces annonene in accordance with the pathway proposed by Chen et al. (Chen *et al*., 2017). This difference could have been due to the use of different cytochrome P450 reductases used to support P450 activity in yeast, or due to differences in the assay environments, particularly conducting the P450 assays without a co-expressed class I diTPS. The coupled *in vivo* assay of SdANS (CYP76AH39) with SdHDAS (CYP728D26) showed the oxidation of annonene to hardwickiic acid as has been postulated for salvinorin A biosynthesis (Chen *et al*., 2017). Although it is possible that crotonolide G is also an intermediate in salvinorin A biosynthesis, given the reported presence of hardwickiic acid in *S. divinorum* (Bigham *et al*., 2003; Chen *et al*., 2017) and the lack of reports of clerodanes with a dihydrofuran ring, to date, we suggest that furanoclerodane formation in *S. divinorum* relies predominantly on the annonene synthase activity of CYP76AH39.

The presence of semi-neofunctionalized enzymes in *S.splendens*, capable of catalysing reactions in both clerodane and other diterpenoid pathways, suggests that the rise of furanoclerodane pathway in Salvia is a relatively recent event. This phylogenetic development likely preceded the diversification of multiple clerodane-rich species. The enzymatic activity of SsHDAS, SsHTAS, and SdHDAS highlights their key roles in furanoclerodane biosynthesis in the subgenus Calosphace. These cytochrome P450 enzymes belong to clan 85 and syntenic analysis showed the presence of multiple gene copies across different Lamiaceae species and lineages. Their activity in the furanoclerodane biosynthetic pathway might have been a prelude for driving the chemical diversity in the family Lamiaceae (Bathe and Tissier, 2019; Consortium, 2018).

In earlier work, we showed that class I clerodane synthases have neofunctionalized following duplication of genes encoding enzymes located in a ferruginol-related BGC conserved in the family Lamiaceae (Li *et al*., 2023). The emergence of annonene synthase was the result of neofunctionalization of a cytochrome P450 enzyme with ferruginol synthase activity, most likely following WGD (Figure 6). Our findings support and expand further the hypothesis that BGCs can serve as ‘‘metabolic toolboxes’’ providing new biocatalysts for evolving metabolic pathways through gene duplication. Cytochrome P450 enzymes play an important role in the enormous chemo-diversity observed in diterpenoid metabolism of plants from the Lamiaceae family and the discovery of the roles of HDAS and HTAS in furanoclerodanes biosynthesis provides new examples of P450 biocatalysts in Lamiaceae diterpenoid metabolism.

In conclusion, we employed a combination of bioinformatic analysis tools and discovered three new cytochrome P450 enzymes acting in the biosynthesis of clerodanes anonnene, hardwickiic acid and hautriwaic acid in *S. splendens*. We also found the orthologues of SsANS and SsHDAS in *S. divinorum* which had the same catalytic activities, supporting the idea that *S. splendens* can serve as an excellent model for studying furanoclerodane in Salvia and that microevolutionary genomics can facilitate future gene discovery in the biosynthesis of high value clerodanes such as salvinorin A in the genus Salvia.

## Materials and Methods

### Materials

#### Plant material and chemicals

Seeds of *Salvia splendens* Ker-Gawler cv. “Olympic flame” were provided by Prof. Ai-Xiang Dong at Beijing Forestry University. Whole plant tissues including flowers, leaves, stems and roots from flowering *S. splendens* were snap-frozen in liquid nitrogen. The frozen tissue was lysed using a tissue lyser, and RNA was extracted using RNeasy Plant Mini Kit (Qiagen, Germany). The extracted RNA was purified with TURBO DNA-free kit (ThermoFisher) to remove gDNA. Subsequently, the RNA was reverse-transcribed into cDNA using SuperScript IV VILO Master Mix (ThermoFisher, USA). The chemicals (-)-hardwickiic acid and hautriwaic acid were purchased from Biobiopha as reference standards.

## Methods

### Generation of self-organizing map

We obtained two sets of RNA-seq data from NCBI SRA project PRJNA422035. Each set consisted of triplicates of RNA-seq data obtained from different tissues (roots, stems, leaves, calyx and corolla). These RNA-seq data were aligned to the scaffold-level *S. splendens* reference genome using Hisat2 v2.1.0 (Kim et al., 2019), and converted to fpkm values using Samtools v1.11 (Danecek et al., 2021) and Cuffdiff (Trapnell et al., 2012). The generation of self-organizing map was in accordance with the method described by Payne et al. (2017), using Kohonen package 3.0.11 (Wehrens and Buydens, 2007). To allow a more extensive search of genes, the map size was determined to allow at least 70 genes cluster in every node. Multiple seed numbers were evaluated to explore the various clustering patterns, and the representative map was selected based on its inclusion of multiple P450 genes that ranked highly in the mutual ranking profile with *SsKPS* (*SspdiTPS2.1*) and were located within the same node as *SsKPS*.

### Mutual rank-based co-expression analysis

The fpkm values obtained above served as input for the MutRank program, which calculated the average ranking of genes based on their correlation with a reference gene (Obayashi and Kinoshita, 2009; Poretsky and Huffaker, 2020).

### Syntenic analysis of CYP76 and CYP716B genes

Genomic data of *Callicarpa americana* (Hamilton et al., 2020), *Scutellaria barbata* (Li *et al*., 2023), *Salvia splendens* (Jia *et al*., 2021), *Salvia miltiorrhiza* (Song et al., 2020), *Salvia bowleyana* (Zheng et al., 2021), *Salvia rosmarinus* (Han et al., 2023) and *Salvia hispanica* (Wang *et al*., 2022a) were downloaded from online depository. MCScan from JCVI package (Tang et al., 2008) was employed to search for interspecies collinear blocks using default settings. The color-coding of cytochrome P450 genes and related diterpene synthases was determined according to the identified gene functions (Li *et al*., 2023), Refseq function annotation or the *in vivo* assay results of this study.

### *De novo* assembly of *S. divinorum* trichome transcriptome

Illumina RNA-seq reads of *S. divinorum* peltate trichome data was downloaded from NCBI (SRR3716680) and assembled into transcripts with Trinity 2.9.1 (Grabherr et al., 2011). Homologous genes of *SspdiTPS1.3*, *SspdiTS1.5*, *SsANS* and *SsHDAS* in *S. divinorum* were identified by blast search into the trichome transcriptome.

### Phylogeny location of *S. divinorum* class I diTPS and CYP genes

SdKSL1, SdKSL2 and SdKSL3 (NCBI accession: KX268507-KX268509)(Chen *et al*., 2017) and (NCBI accession: KY057342-KY057344)(Pelot *et al*., 2017) were added to the phylogenetic tree modified from Figure 5A in Li *et al*. (2023), using MAFFT (Katoh and Standley, 2013) for alignment and iqtree-1.6.10 (Nguyen et al., 2015) to build the tree. The phylogenetic tree for the two studied CYP families was constructed with the same pipeline. The sequences included were derived from genes syntenic to those characterized CYP76AH genes or HDAS gene in *S. splendens*, as well as from unigenes found in the trichome transcriptome of *S. divinorum*.

### Species divergence time estimation

Orthofinder 2.5.4 (Emms and Kelly, 2019) was applied to construct a species phylogenetic tree. The genome of *Oryza sativa* Japonica (IRGSP1.0, EnsemblPlants), *Solanum lycopersicum* (SL3.0, EnsemblPlants), *Antirrhinum majus* (doi: 10.1038/s41477-018-0349-9), *Callicarpa americana* (NCBI: PRJNA529675), *Scutellaria barbata* (NCBI: PRJNA649842), *Salvia rosmarinus* (doi: 10.6084/m9.figshare.21443223.v1), *Salvia bowleyana* (National Genomics Data Center: PRJCA003734), *Salvia miltiorrhiza* (National Genomics Data Center: PRJCA003150), *Salvia hispanica* (China National GeneBank DataBase: CNA0047366) and *Salvia splendens* (doi: 10.1093/gigascience/giy068) were used for speciation time estimation. The longest protein sequence isoform from each genome were supplied to Orthofinder for producing a species tree. Speciation time estimation was carried out with mcmctree from PAML (Yang, 2007), using the phylogenetic tree generated from Orthofinder and the species divergence times were estimated by TimeTree (Kumar et al., 2017) as calibration points (*O. sativa* and *S. lycopersicum*, *S. lycopersicum* and *A. majus*).

### Bayesian analysis on syntenic cytochrome P450s from CYP76AH and CYP716B/CYP728D subfamilies

The Bayesian phylogenetic trees of proteins encoded by syntenic *CYP76AH* and *CYP716B* genes were constructed with Beast2 (Bouckaert et al., 2014). Translated *CYP76AH* or *CYP716B* gene sequences were first aligned using MAFFT (L-INS-i) (Katoh and Standley, 2013), then converted to codon alignment with PAL2NAL (Suyama et al., 2006). The estimation of gene divergence time was carried out with the following priors: HKY85 substitution model, empirical frequencies, strict molecular clock and a Calibrated Yule Model. The estimated species divergence time obtained from Orthofinder and PAML served as the calibration points (Bouckaert *et al*., 2014). The MCMC chain length for posterior probability was set at 20,000,000. A 10% burn-in was applied to the tree using treeannotator, and plotted using Figtree.

### cDNA cloning

The coding sequence of P450 genes were isolated using primers listed in Supplementary file Table S2. Initial amplification of candidate genes were either performed with KOD plus Neo (TOYOBO, Japan) or Phusion Plus PCR Master Mix (ThermoFisher, USA). PCR products were isolated and ligated onto pESC vectors using In-Fusion cloning kit (TakaraBio, Japan), which were subsequently transformed into DH5α competent cells. Transformed colonies were screened on carbenicillin selective media (50 mg L^−1^) and colonies that showed the correct insertion size in colony PCR were picked for overnight incubation in 5 mL carbenicillin selective liquid Luria-Bertani medium. Plasmids were extracted with TIANprep Mini Plasmid Kit (Tiangen) and sent for sequencing to confirm sequence identity.

pESC vectors have two multiple cloning sites, Gal1 and Gal10. All Gal1 sites were digested with BamHI-SalI and all Gal10 sites were digested with SpeI, hence for genes inserted at Gal1 site the primers added a BamHI site at the 5’ end and a SalI site at the 3’ site. For genes inserted at the Gal10 site, the primers added an SpeI site on both 5’ and 3’ ends.

### Yeast vector construction

#### pESC-His vector for kolavenol production

To introduce kolavenol pathway into yeast, a pESC-His vector carrying a GGPP synthase gene and a fused diterpene synthase gene was constructed. Erg20 is a farnesyl pyrophosphate synthase endogenous to yeast, however its F96C mutant functions as a GGPP synthase (Ignea et al., 2015). *ERG20* gene was cloned from yeast cDNA and was subjected to site-directed mutagenesis using primers listed in Table S2 to produce *ERG20*::F96C. The mutated *ERG20*::F96C was introduced into Gal1 site of pESC-His vector. Kolavenol production was achieved by introducing a kolavenol synthase (SspdiTPS1.5Δ50) fused a kolavnyl pyrophosphate synthase (SspdiTPS2.1). This approach of fusing a class I diTPS with a class II diTPS has been shown to be sufficient for producing the desired diterpenoid scaffold in following expression in yeast (Zhou *et al*., 2012). In our case, *SspdiTPS1.5*Δ*50* and *SspdiTPS2.1* were linked by a GGGS linker (5’ GGT GGT GGT TCT 3’), with the stop codon of *SspdiTPS1.5*Δ*50* replaced by the GGGS linker to allow a complete translation of the fused protein. A start codon was also added in front of *SspdiTPS1.5*Δ*50*.

#### pESC-His vector for miltiradiene production

The introduction of miltiradiene pathway into yeast followed the same procedure as described above for kolavenol pathway. The gene inserted into Gal10 site was replaced by a fusion of truncated miltiradiene synthase SspdiTPS1.3Δ61 with full length copalyl diphosphate synthase SspdiTPS2.4.

#### pESC-Leu vector for CYP76 screening

The full functionalization of CYP required the presence of cytochrome P450 oxidoreductase (CPR), therefore *CrCPR2* from *Catharanthus roseus* was introduced into the Gal1 site of pESC-Leu vector (Meijer et al., 1993). CYP76 genes were inserted into Gal10 site.

#### pESC-Ura for CYP716B/CYP728 screening

In the initial screening stage, all three CYP716B genes were ligated onto the Gal1 site of the pESC-Ura vector. Later when Saspl_010504 was identified as the hardwickiic acid synthase, *Saspl_010503* and *Saspl_010505* was subcloned onto the Gal10 site of pESC-Ura vector harboring HDAS in the Gal1 site.

### Heterologous expression in yeast and *in vitro* activity assay

Yeast strain AM119 was used as the host strain for CYP screening (29). Depending on the screening purposes, different combinations of pESC vectors were co-transformed into the host strain (See Table S3 for details). For heterologous expression, single yeast colonies were picked from fresh transformation and incubated in 2% glucose supplied SD dropout media for 12 h. The seed cultures were then added into 20 mL fresh glucose-supplied SD dropout media, grew to OD600=0.6-0.8, washed and pelleted before adding into 20 mL SD dropout media supplied with 2% galactose. Induced cultures were incubated on a 30 °C shaker for 48 h. The cultures were then centrifuged, and the pellets were lysed in tissue-lyser. The supernatant and lysed pellets were pooled back together and extracted with equal volume of ethyl acetate three times. Ethyl acetate extracts were fully evaporated on a rotary evaporator, then resuspended in 50 µL of hexane for GC-MS analysis or 150 µL of methanol for LC-MS analysis.

### Protein expression and purification

LB media containing selective antibiotics were inoculated with selected recombinant *E. coli* colonies and cultured overnight at 220 rpm, 37 °C. This overnight culture was used to inoculate Terrific-Broth media containing the same selective antibiotics at 1:50 dilution. The TB culture was incubated at 37 °C and shaken at 220 rpm until the optical density at 600 nm reached 0.8-1.0. Cultures were cooled to 16 °C for 30 min and induced with 1 mM isopropylthio-β –galactoside. Cultures were incubated at 16 °C for a further 16 h before centrifugation. The pellets were resuspended in chilled lysis buffer (50 mM tris-HCl buffer, 50 mM glycine, 5% v/v glycerol, 0.5 M NaCl, 20 mM imidazole, 0.2 mg mL−1 lysozyme and 1 mM phenylmethylsulfonyl fluoride or Complete Protease Inhibitor Cocktail [Roche], pH=8) and kept on ice for 30 min. The lysate was disrupted by sonication (Scientz JY92-IIN) and centrifuged at 4 °C (35000 rpm, 20 min). A slurry of 200-250 μL Ni NTA beads (Smart LifeSciences) was added into the collected supernatants to pull down the His-tagged protein. After gently shaking the mixture on a rocking platform at 4 °C for 1.5 h, the mixture was centrifuged and washed with ice cold lysis buffer (that did not contain protease inhibitor or lysozyme) twice. Proteins were eluted with 600 μL elution buffer (50 mM tris-HCl buffer pH=8, 50 mM glycine, 5% v/v glycerol, 0.5 M NaCl, 0.5 M imidazole) and the eluate were applied to a 4 mL Amicon Centrifugal 30k NMWL tube for dialysis, where the buffer content was exchanged to phosphate-buffered saline (0.137 M NaCl, 0.0027 M KCl, 0.01 M phosphate buffer, pH = 7.4). Protein concentration was measured by the A280 using Nanodrop and diluted to approximately 1 mg mL-1 to avoid precipitation. Purified enzymes were divided into aliquots, snap frozen in liquid N2 and stored under −80 °C.

### Class I diterpene synthases *in vitro* enzyme assays

Enzyme assay buffer was consisted of 50 mM tris-HCl (pH = 7.2), 100 mM KCl, 7.5 mM MgCl_2_, 5mM DTT and 5% glycerol, and GGPP was added to 100 µM in 250 uL assays. Around 100 µg class II diterpene synthase SsKPS alone or in combination with 80 µg class I diterpene synthases SdKSL1 and SdKSL2 were used in the reaction system. Assays were incubated at 37 °C overnight in the dark. For the SsKPS enzyme assay positive control, the assay mixture was further incubated with 2 μL alkaline phosphatase (CIAP Takara Bio) for 2 h. Enzyme products were extracted with hexane for three times, and then dried and resuspended with 50 uL hexane before GC-QTOF analysis.

### GC-MS analysis

Ethyl acetate extracts of yeast products were analysed using an Agilent DB-5HT (30 m × 0.25 mm × 0.1 μm) column on a GC7890B-MS7200B QTOF. Samples were injected in 2 μL volume in split mode (split ratio 5:1) at an inlet temperature of 250°C. The gradient of the GC program was as follows: hold at 50°C for 2 min; increase to 110°C at 4°C min^-1^; increase to 250 °C at 8°C min^-1^; increase to 310 °C at 10 °C min^-1^ and hold for 5 min. The temperature of the mass spectrometer for ion source was set to 230 °C and spectra were recorded from m/z 50 to m/z 350.

### LC-MS analysis

Ethyl acetate extracts of yeast products were dried with rotary evaporator, and then dissolved in methanol and measured by LC-MS (Q-Exactive plus) using an Agilent BEH C18 (100×3.1 mm) column with 2.6 μm particle size. The aqueous eluent phase A was 2 mM ammonium formate + 0.01% formic acid, and the organic eluent phase B was acetonitrile. The conditions were set as follows: increase the concentration of B from 40% to 99% from 0 to 10 min, and hold at 99% for 2 min.

### qRT-PCR

The tissues (roots, stems, flowers, leaves) of Salvia splendens were collected at the flowering stage and each tissue was extracted with three biological replicates. The RNA was extracted using the CTAB method. The subsequent experiments were performed with TB Green® Premix DimerEraser (Perfect Real Time, Takara) reagent on an AB StepOne Plus system, with SspqACTIN1 used as the reference gene. The qRT-PCR conditions were: Initial denaturation at 95°C for 30 s, followed by 40 cycles of amplifications as follow: 95°C for 3 s, 55°C for 30 s, 72°C for 30 s. Gene expression was calculated as described in our previous work (25). The primers for qRT-PCR are shown in Table S2.

### Salvia splendens tissue extracts

The tissues (root, leaf, flower and stem) of *S. splendens* were collected from greenhouse, snap frozen and ground into powder, then transferred into 2 mL tubes and extracted with acetone at room temperature overnight in a shaker at 200 rpm. The tubes were further sonicated for 10 min twice, and centrifuged at 10000 rpm for 10 min. The supernatant was then dried in a rotary evaporator, and re-dissolved in 150 μL methanol before LCMS (Q-Exactive) examination. Clerodanes were tentatively identified through mass spectra based on molecular weight and exact mass.

#### *S. splendens* abaxial peltate trichome extracts

Peltate trichomes from the abaxial side of *S. splendens* leaf (about 50 μm in diameter) were manually collected with blade tip under a dissection microscope. About 100 – 200 trichomes were pooled together and extracted with MeOH. The trichome extracts were evaporated down to 50 μL in volume, and 5 μL of the extract were examined by an Agilent Q-Exactive equipped with an atmospheric pressure chemical isonisation (APCI) source. Metabolite analysis was performed with an Agilent Kinetex C19 column (2.6 μm, 2.1 x 100 mm). The mobile phase with a flow rate of 0.4 mL.min-1 consisted of solvent A (2 mM ammonium formate with 0.1% formic acid) and solvent B (ACN). A binary gradient elution was performed as follows: linear gradient 5-98% B from 0 to 8.0 min, followed by 98% B from 8.0 to 10.0 min, then returned to 5% B at 10.1 min and maintained until 14.0 min for column equilibration. Clerodanes were tentatively identified through mass spectra based on molecular weight and exact mass.

### MALDI-MS imaging analysis

Mature leaves of *S. splendens* were clamped flat between two glass slides and vacuum dried in a desiccator. Optical images of the abaxial side of each leaf were taken using a Canon 5D Mark IV camera with a Canon MP-E 65=mm=f/2.8 1–5x Macro Photo lens (Canon). Leaf samples were covered with 2,5-dihydroxybenzoic acid matrix (DHB, Sigma-Aldrich) using a SunCollect MALDI Sprayer (SunChrome) with a DHB solution of 10=mg=ml^−1^ in 80% methanol to a density of approximately 3=µg=mm^−2^. MALDI imaging was performed with a Synapt G2-Si mass spectrometer with a MALDI source (Waters) equipped with a 2.5=kHz Nd:YAG (neodymium-doped yttrium aluminum garnet) laser operated at 355=nm. Red phosphorous clusters were used for instrument calibration.

The slides with the leaves were fixed in the instrument metal holder and were scanned with a flat-bed scanner (Canon). The images were used to generate pattern files and acquisition methods in the HDImaging software version 1.4 (Waters). An area of approx. 6.5×4 mm was selected and scanned with the 60 µm laser beam with steps of 45 µm. The instrument was set to MALDI–MS-negative sensitivity mode, *m*/*z* 100–1,200, scan time 0.5=s, laser repetition rate 1=kHz, laser energy 220.

The MS raw files were processed in HDI1.4 with the following parameters: detection of the 2,000 most-abundant peaks, *m*/*z* window 0.05, MS resolution 10,000; a list with the masses of interest was loaded as a target mass file. The processed data were loaded into HDI1.4 and normalized by total ion content. The MALDI images were generated using the HotMetal2 colour scale and overlaid with the original optical image to show the location of the MALDI signals.

### Associated Content

**Supporting Information:** the file (PDF) contains supporting Figures S1 to S17, supporting Tables S1 to S3, Legends for Dataset S1, and SI References.

**Dataset:** the file (EXCEL-XLS) contains the data from self-organizing map (SOM) and mutual ranking analysis (MR)

## Acknowledgments

The authors would like to thank the staff and management of the CEMPS Core Facility Center for the excellent support in metabolomics (LC-MS, NMR) and microscopy services, the personnel at the CEMPS glasshouse facilities and Matt Downie (John Innes Centre) for carrying out SEM imaging works. The authors thank Prof A. M. Makris (INAB-CERTH) for providing the AM119 yeast strain.

## Data availability

Gene sequence data for this study have been deposited in the National Center for Biotechnology Information (NCBI) database OR837137 - OR837141.

