## Supplementary material for "Three cytochrome P450 enzymes catalyse the formation of furanoclerodane precursors in Salvia spp": SI

### Supporting Information for
Three cytochrome P450 enzymes catalyse the formation of furanoclerodane precursors in Salvia spp.

Ruoxi Lin#, Haixiu Li#, Yiren Xiao, Zhuo Wang, Licheng Liu, Gerhard Saalbach, Carlo Martins, Cathie Martin, Evangelos C. Tatsis*

Evangelos C. Tatsis


**This PDF file includes:**

Figures S1 to S17
Tables S1 to S3
Legends for Dataset S1
SI References

**Other supporting materials for this manuscript include the following:**

Dataset S1

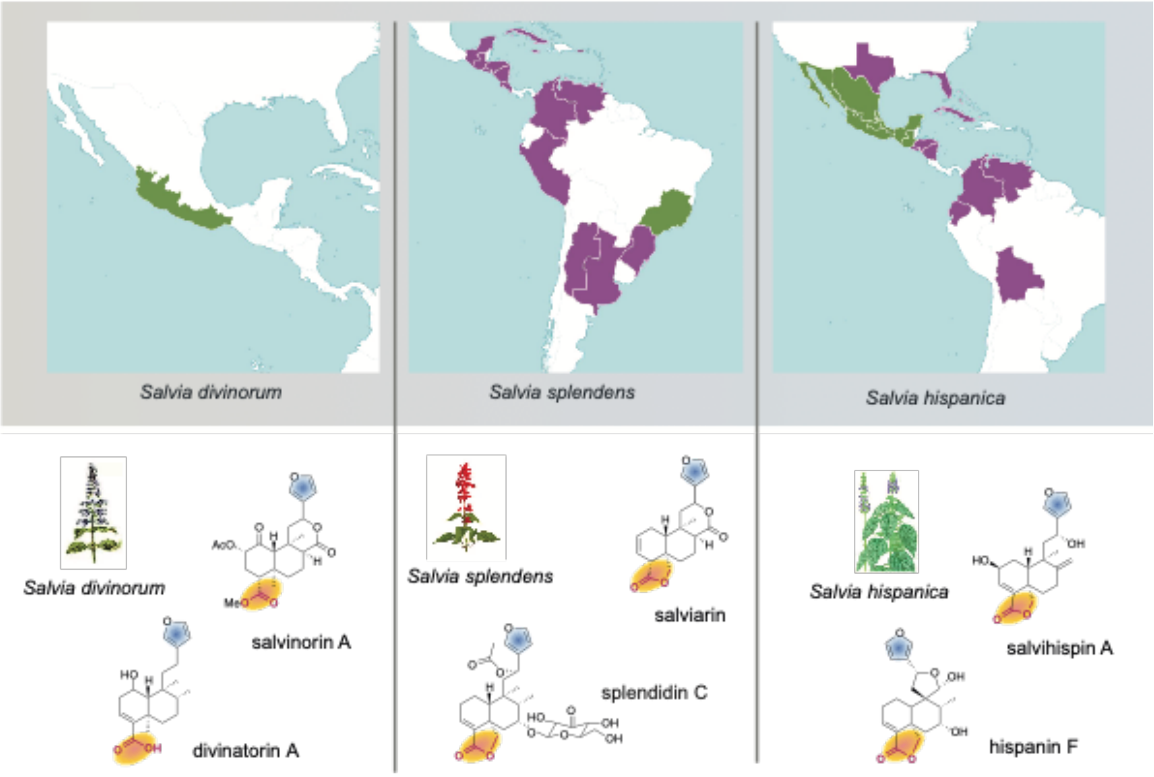


**Fig. S1. Geographical distribution of the three neotropical sages *Salvia divinorum* (magic sage), *Salvia splendens* (scarlet sage) and *Salvia hispanica* (chia) and their clerodane diterpenoids.** The maps were generated by Royal Kew Gardens Plants of the World Online (https://powo.science.kew.org/). Regions labelled in green are the native habitats and regions in purple are the introduced habitats. Examples of furanoclerodanes bearing the furane ring (highlighted in blue) and the oxidized C18, C19 atoms (highlighted in orange) from neotropical sages: the ornamental scarlet sage (*Salvia splendens*), the psychoactive magic sage (*Salvia divinorum*), and the pseudocereal chia (*Salvia hispanica*).

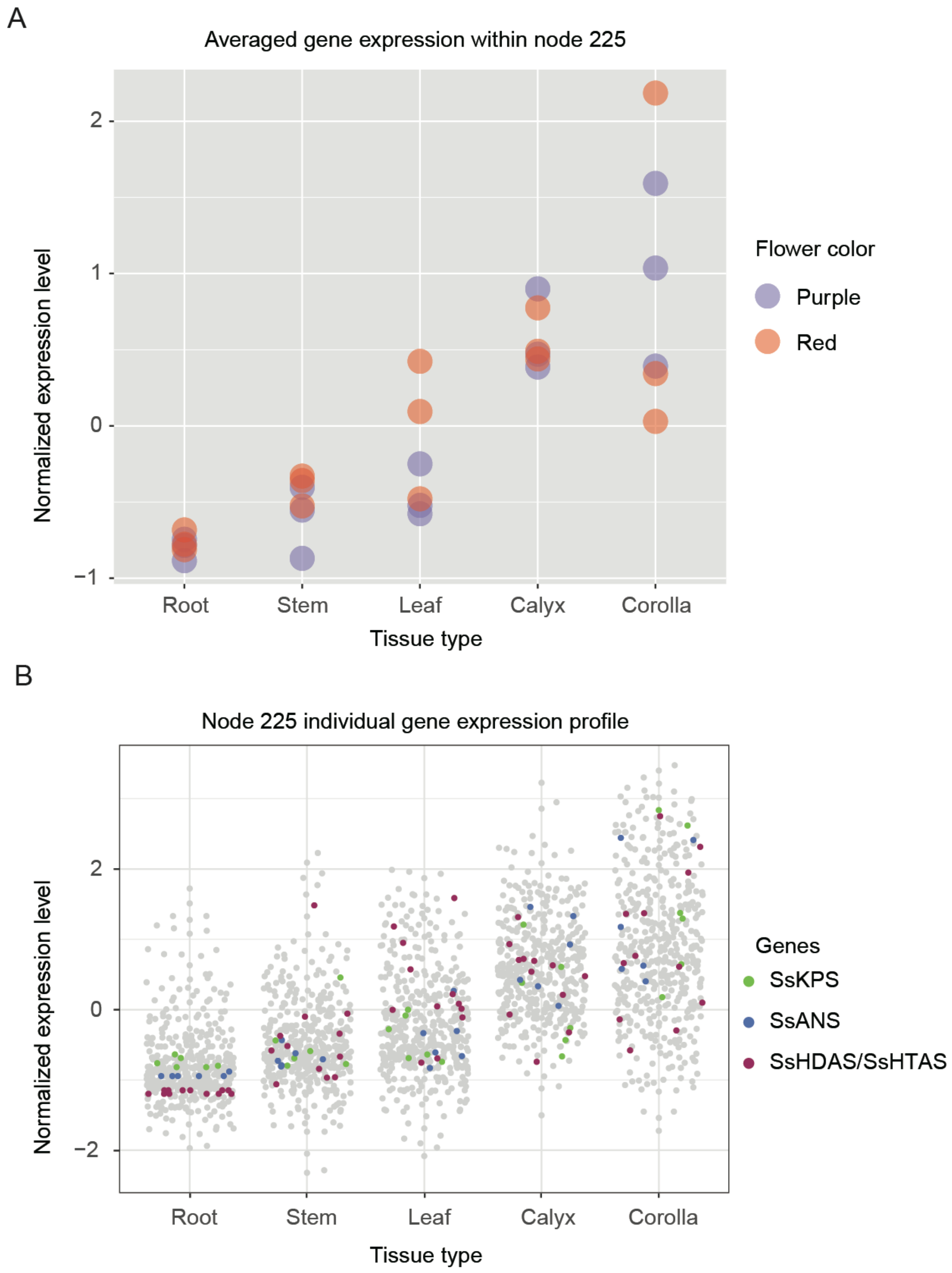


**Fig. S2. *Salvia splendens* gene expression of SOM node 225. A**, averaged gene expression level across tissues. Every circle symbolizes the average FPKM value of all genes found in node

225 within a single sample. The hue of the circle corresponds to the plant phenotype (flower color) that was sampled. There are three biological replicates for each tissue type in both purple flower and red flower phenotypes. **B**, the FPKM value of individual genes across different samples. *SsKPS* (*SspdiTPS2.1*), *SsANS* (CYP76AH), *SsHDAS* and *SsHTAS* (CYP716B/CYP728D genes) are highlighted with corresponding colors.

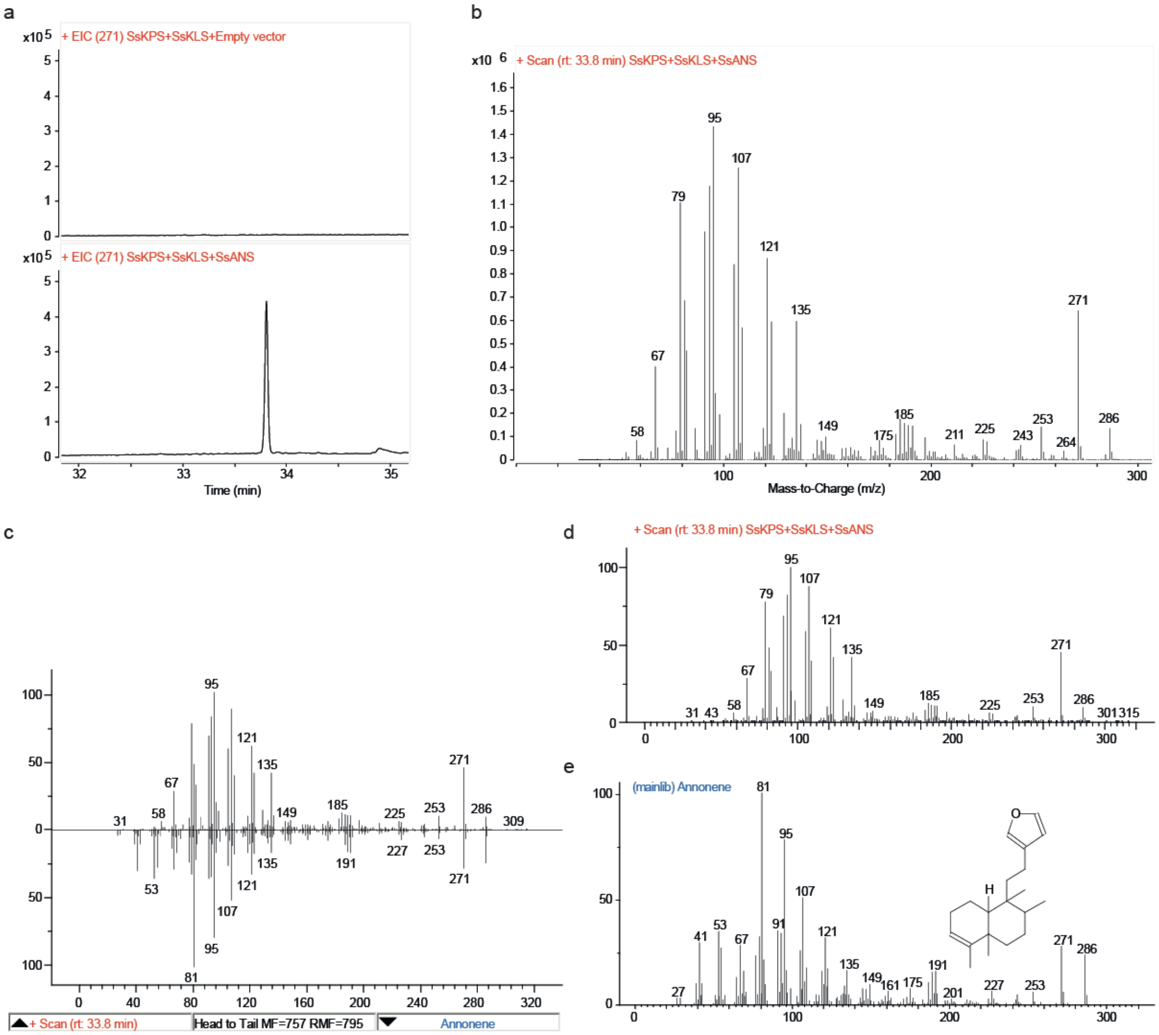


**Fig. S3. GC-MS analysis of SsANS product. a;** GC-MS profile (selected *m/z* 271) of yeast strains expressing GGPPS (Erg20::F96C), SsKPS, SsKLS, CrCPR, empty vector or SsANS. **b;** MS spectrum of peak at 33.8 min. **c;** Comparison of MS spectrum of SsANS product eluted at 33.8 min with the annonene standard (EI MS spectrum from NIST 17 library), and the match probability is 75.2%. **d;** Submitted MS spectrum of SsANS product eluted at a33.8 min, **e**; EI MS spectrum of annonene as it is deposited in NIST17 library.

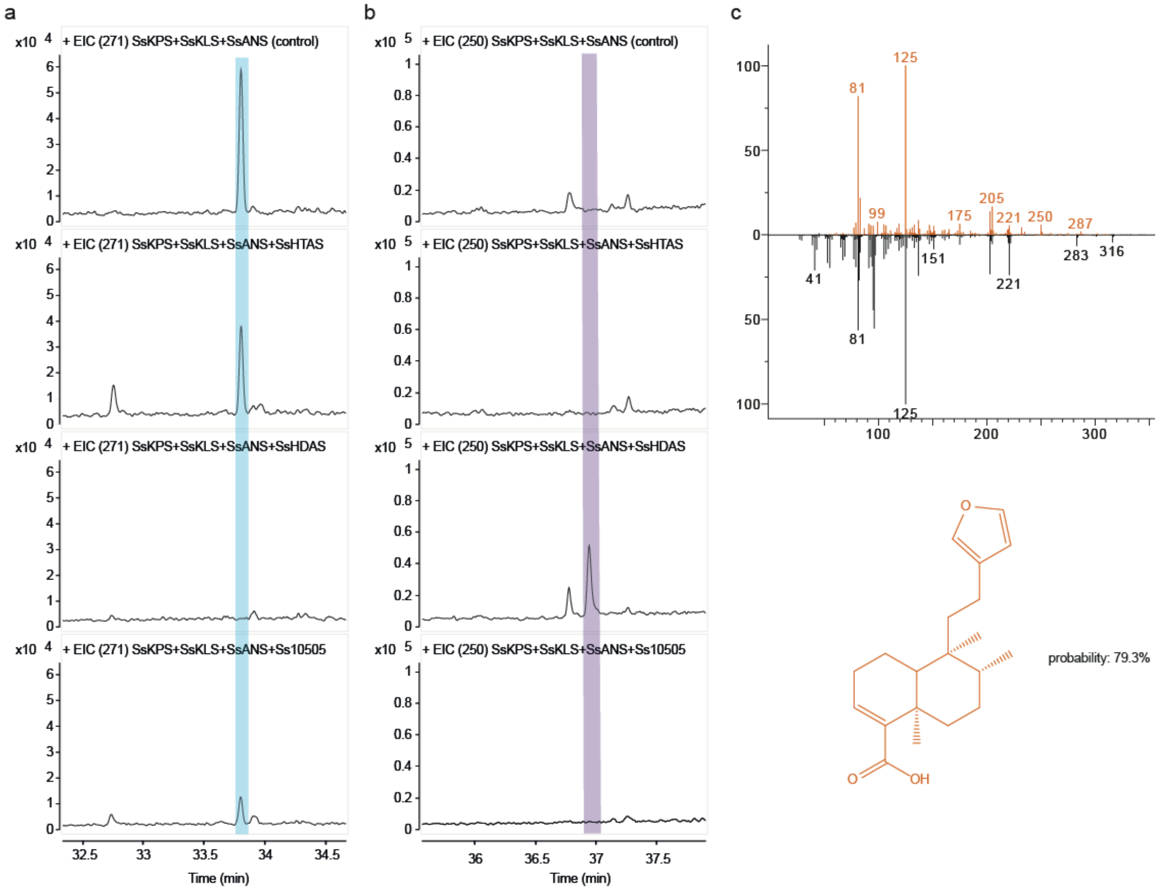


**Fig. S4. GC-MS analysis of HDAS product. a,b;** GC-MS profile (selected m/z 271 and 250) of yeast strains expressing GGPPS (Erg20::F96C), SsKPS, SsKLS, CrCPR, SsANS, empty vector or different CYP716B/CYP728 family proteins. **c;** Comparison of MS spectrum of SsHDAS product with the hardwickiic acid (EI MS spectrum from NIST 17 library), and the match probability is 79.3%.

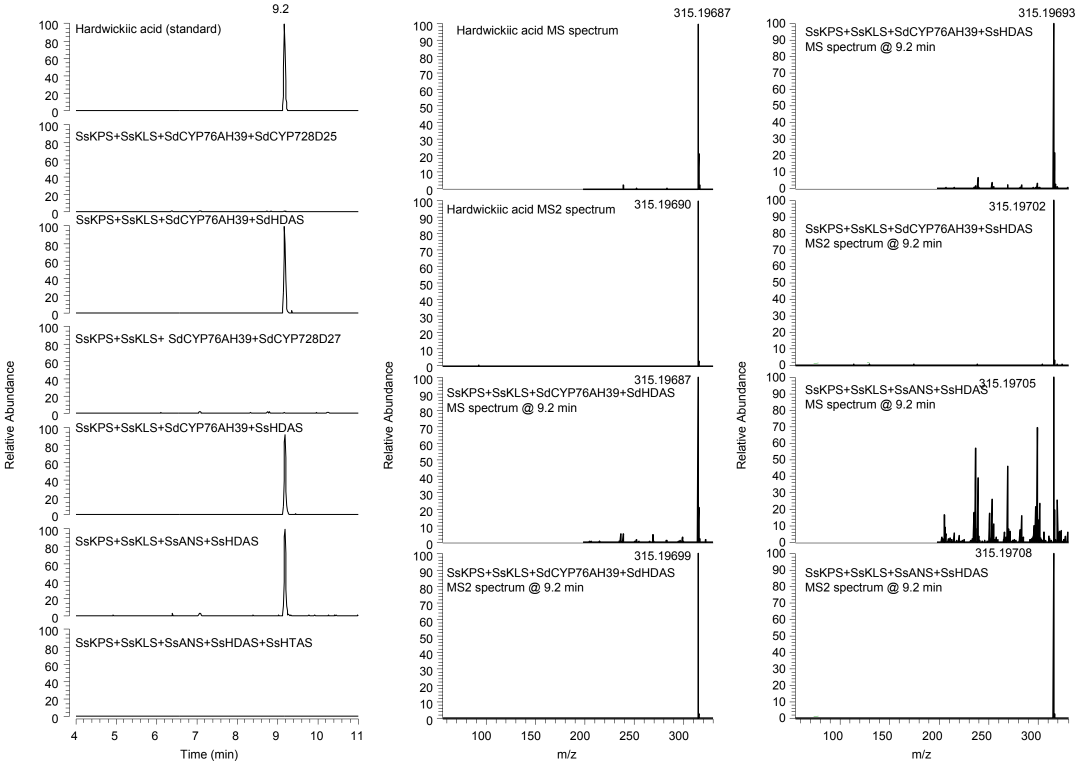


**Fig. S5. LC-MS analysis of HDAS product**. **A**, LC-MS chromatogram depicting different assemblies of clerodane biosynthesis genes compared with the hardwickiic acid standard. The sole combination of SsKPS, SsKLS, SsANS or SdANS, and SsHDAS or SsHDAS yields hardwickiic acid; when incorporating HTAS into the ensemble, the peak corresponding to hardwickiic acid diminishes, implying the role of HTAS in the modulation of hardwickiic acid. **B**, MS and $MS^2$ profile of the hardwickiic acid standard (upper panels) compared to the same profiles of the peak produced by the assembly of SsKPS, SsKLS, SsANS and SsHDAS or SdHDAS (bottom panels).

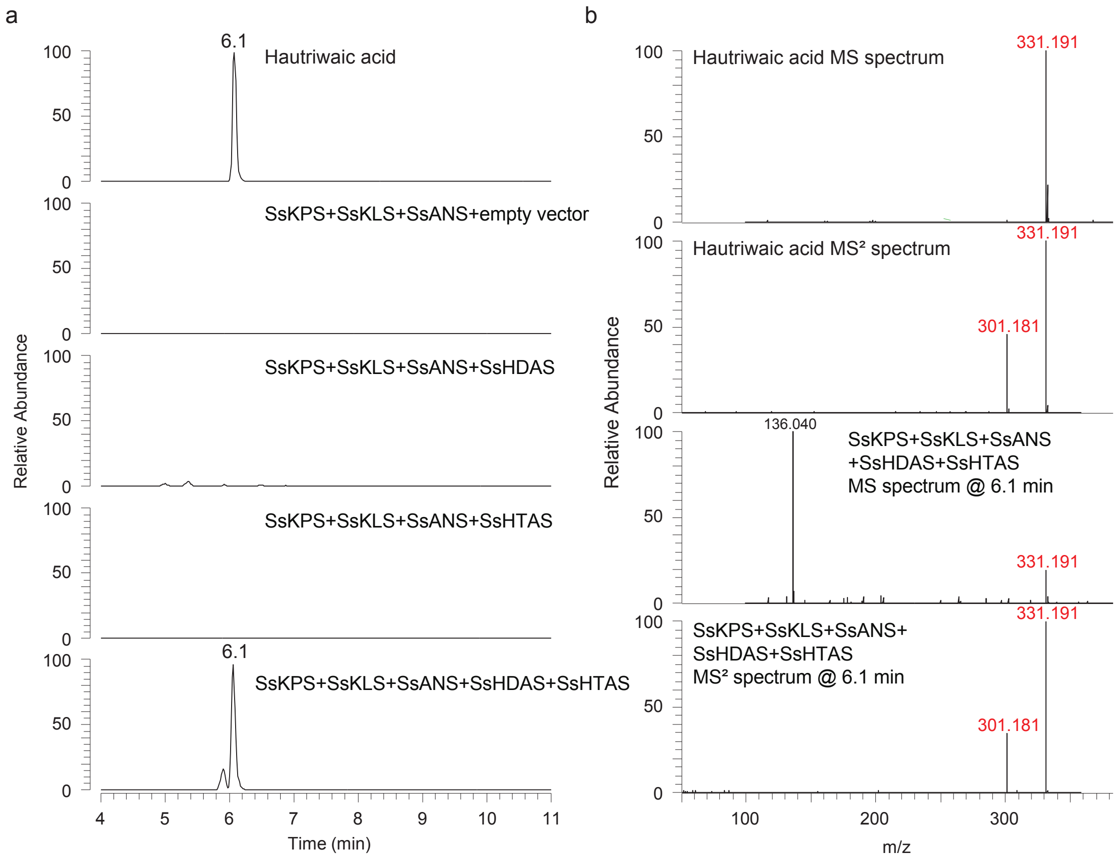


**Fig. S6. LC-MS analysis of HTAS product. a**, LC-MS chromatogram illustrating various combinations of clerodane biosynthesis genes alongside the hautriwaic acid standard. The co-expression of SsKPS, SsKLS, SsANS, SsHDAS, and SsHTAS results in the emergence of a new peak at 6.1 minutes, corresponding to the elution time of the hautriwaic acid standard. **b**, Upper panels: MS and $MS^2$ profile of the hautriwaic acid standard; bottom panels: corresponding profiles of the new peak from the assembly of SsKPS, SsKLS, SsANS, SsHDAS and SsHTAS.

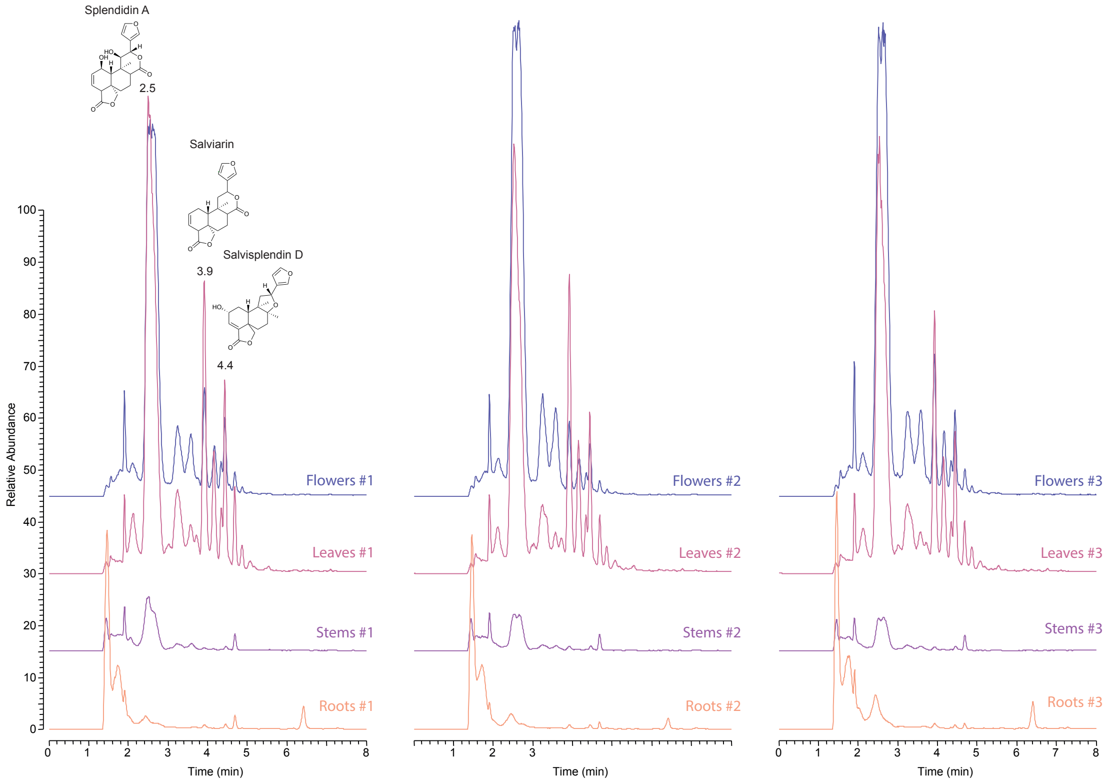


**Fig. S7. LCMS analysis of extracts from different *S. splendens* tissues (flowers, leaves, stems, roots).** Identification of splendidin A, salviarin and salviplendin D was based on molecular weight and exact mass.

A

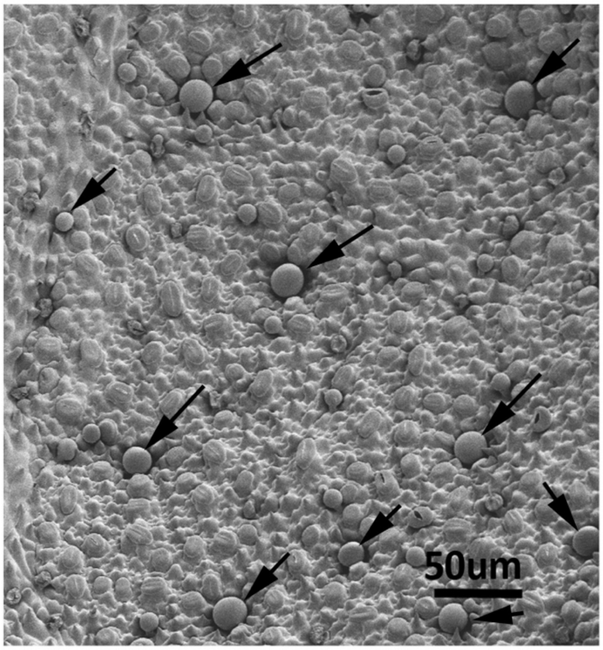


B

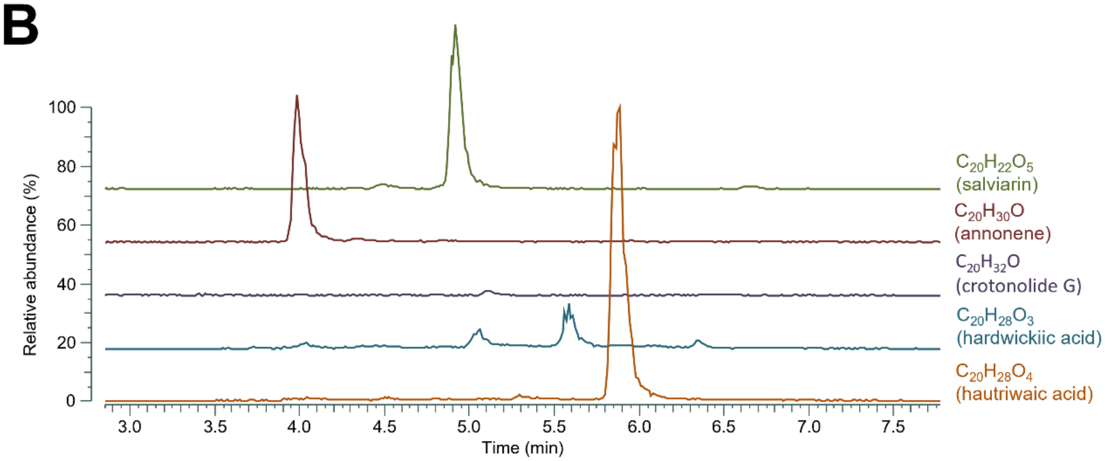


**Fig. S8. Morphology and metabolite content analysis for *S. splendens* leaf abaxial peltate trichomes. A**, SEM of abaxial surface of leaf of *S. splendens.* Black arrows indicate medium sized peltate trichomes picked for LC-MS analysis of metabolite content. **B**, Identification of salviarin, annonene, hardwickiic acid and hautriwaic acid was based on molecular weight and exact mass.

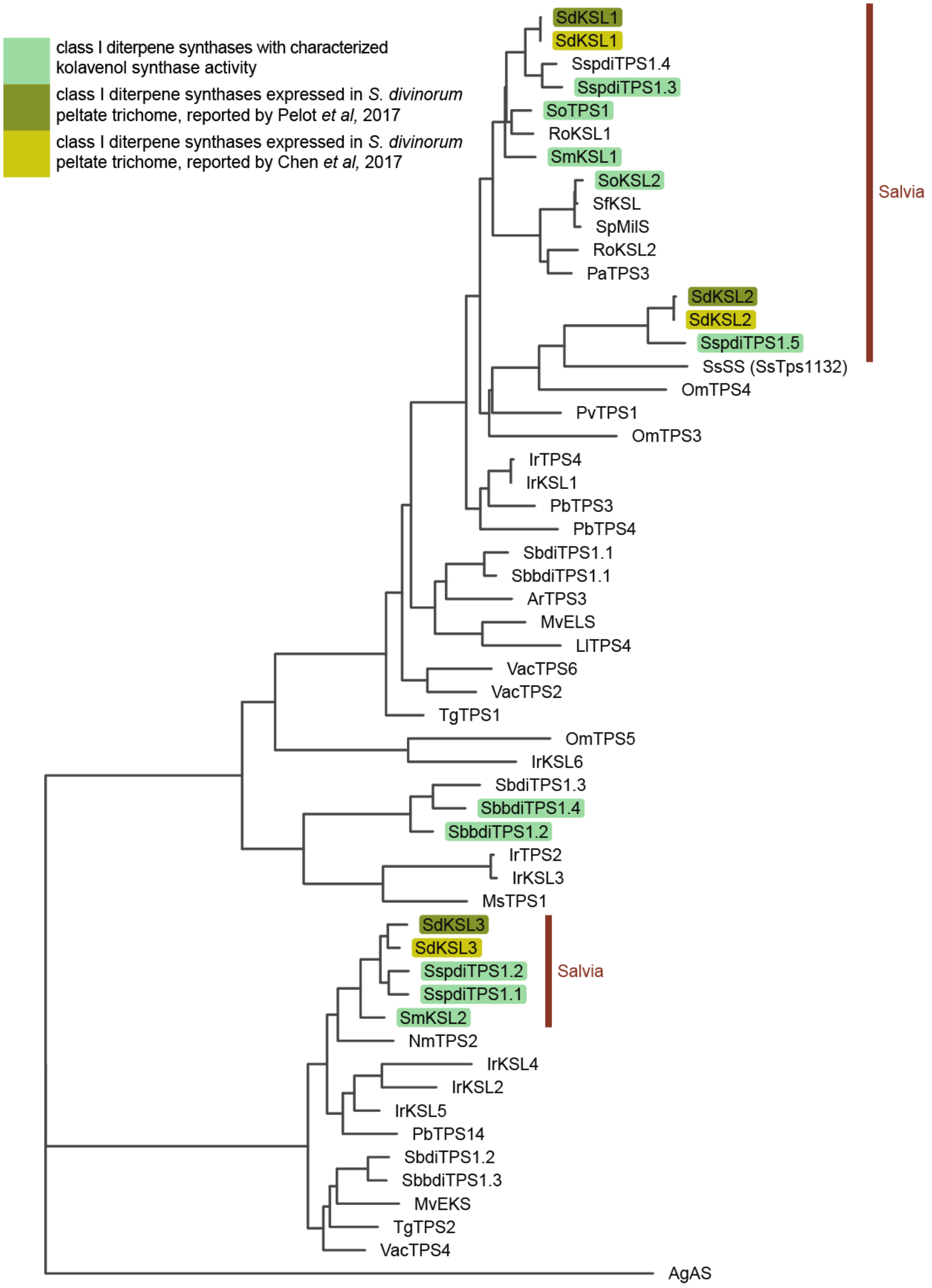


**Fig. S9. Phylogenetic tree of class I diterpene synthases in Lamiaceae.** Bright green highlights the characterized class I diterpene synthases with kolavenol synthase activity. Olive green and mustard green marks the class I diterpene synthases (diTPSs) cloned by peltate trichomes of *S.*

*divinorum* by Pelot et al 2017(Pelot et al., 2017) and Chen et al 2017 (Chen et al., 2017), respectively. Dark brown line indicates the genes belonging to Salvia genus.

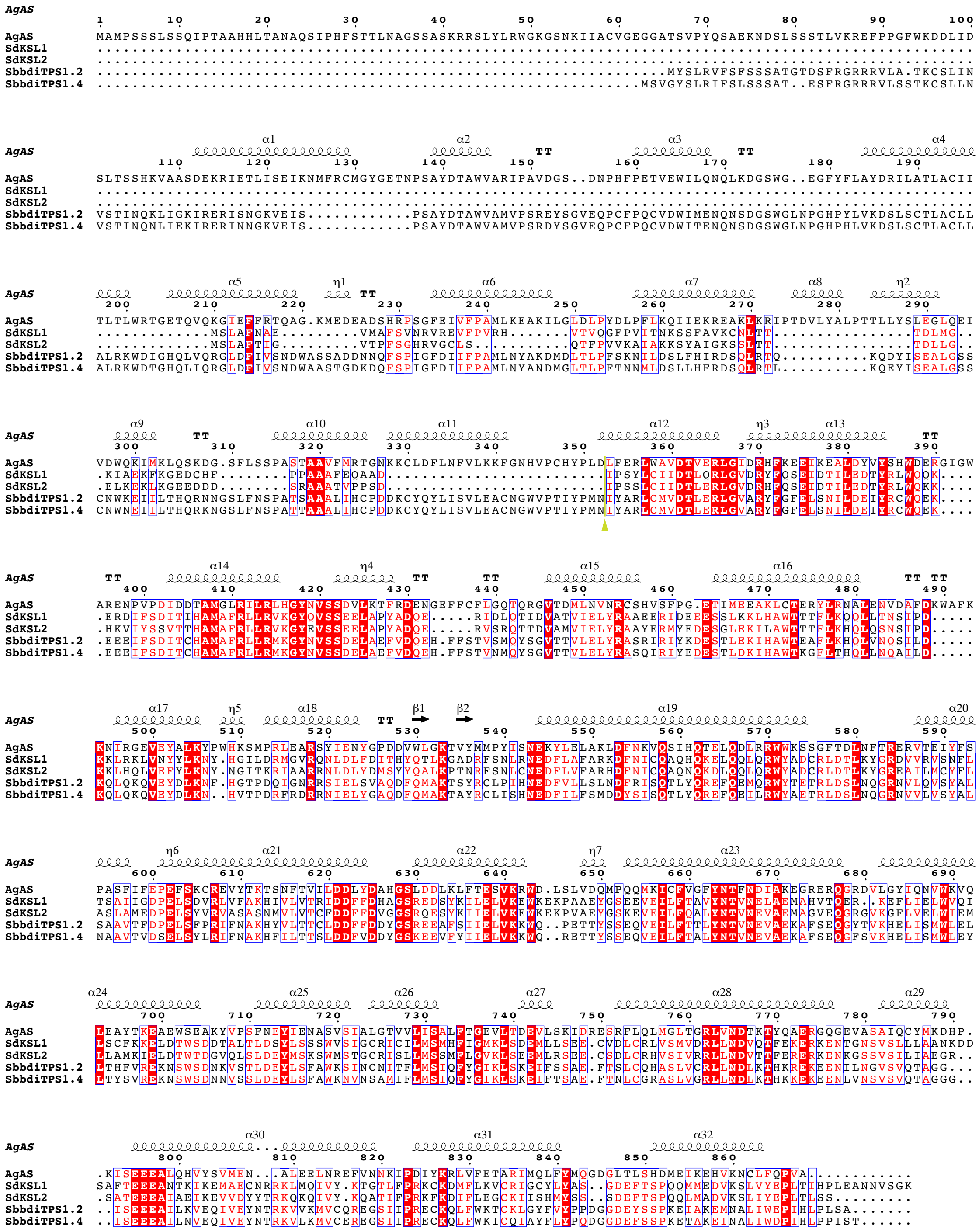

**Fig. S10. Alignment of class I diterpene synthases in family Lamiaceae with kolavenol synthase activity from *Salvia divinorum* (SdKSL1, SdKSL2) (Chen *et al.*, 2017; Pelot *et al.*, 2017) and *Scutellaria barbata* (SbbdiTPS1.2, SbbdiTPS1.4) (Li et al., 2023).** The green arrow and line indicate the beginning of truncated site.

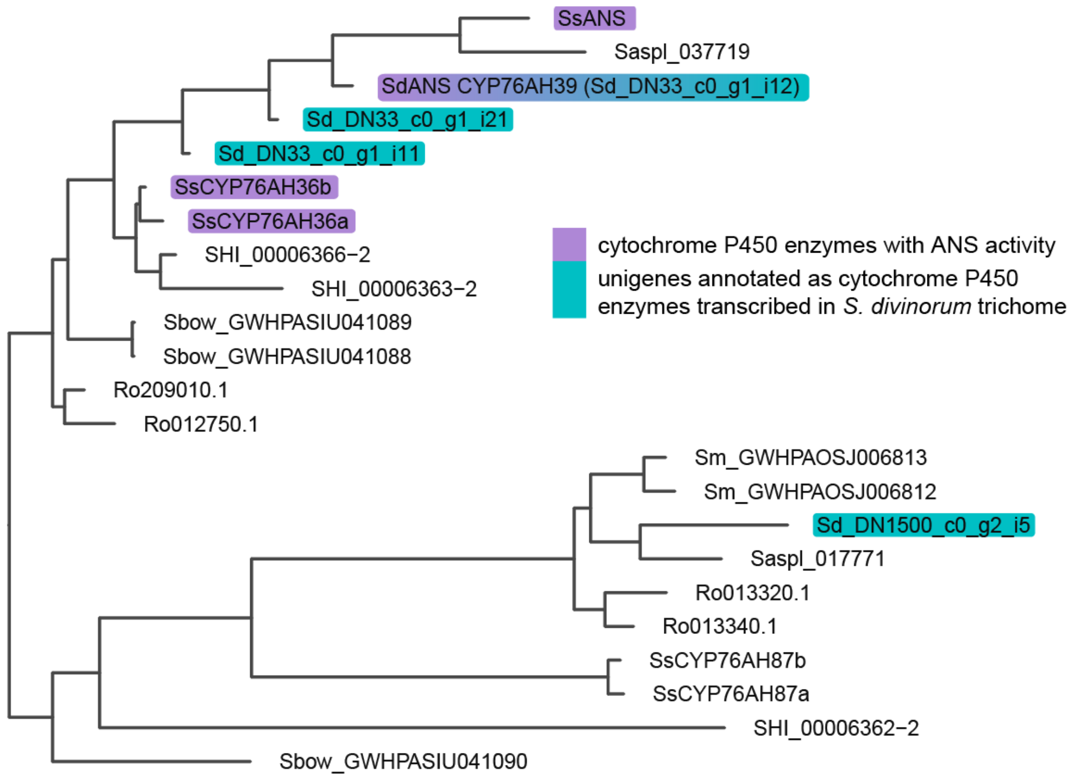


**Fig. S11. Phylogenetic tree of CYP76AH in Salvia genus.** Characterized cytochrome P450 enzymes with annonene synthase activity are highlighted in purple. Their homologous unigenes obtained from *S. divinorum* trichomes EST data are highlighted in teal.

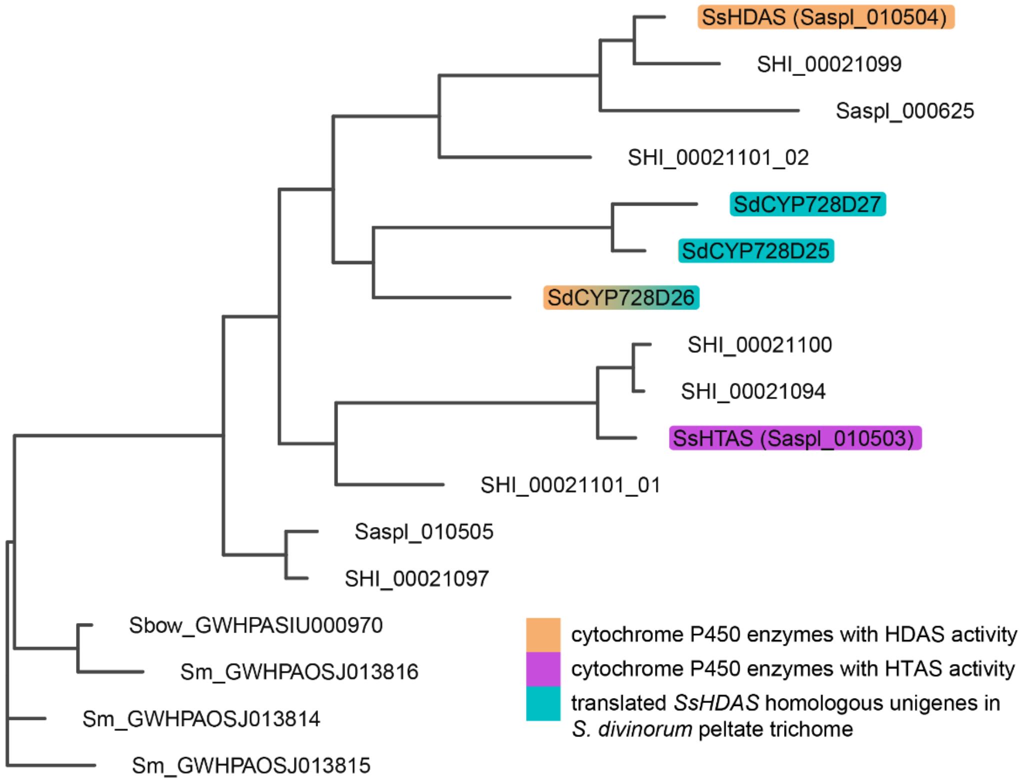


**Fig. S12. Phylogenetic tree of CYP716B/CYP728D in Salvia genus.** Characterized cytochrome P450 enzymes with hardwickiic acid synthase (HDAS) activity, and hautriwaic acid synthase (HTAS) activity, highlighted in orange and violet, respectively. The homologous cytochrome P450 enzymes from *S. divinorum* are highlighted in teal. Shi, Sbow, Sm abbreviations are referred to homologous enzymes from *Salvia hispanica, Salvia bowelyana,* and *Salvia miltiorhiza.*

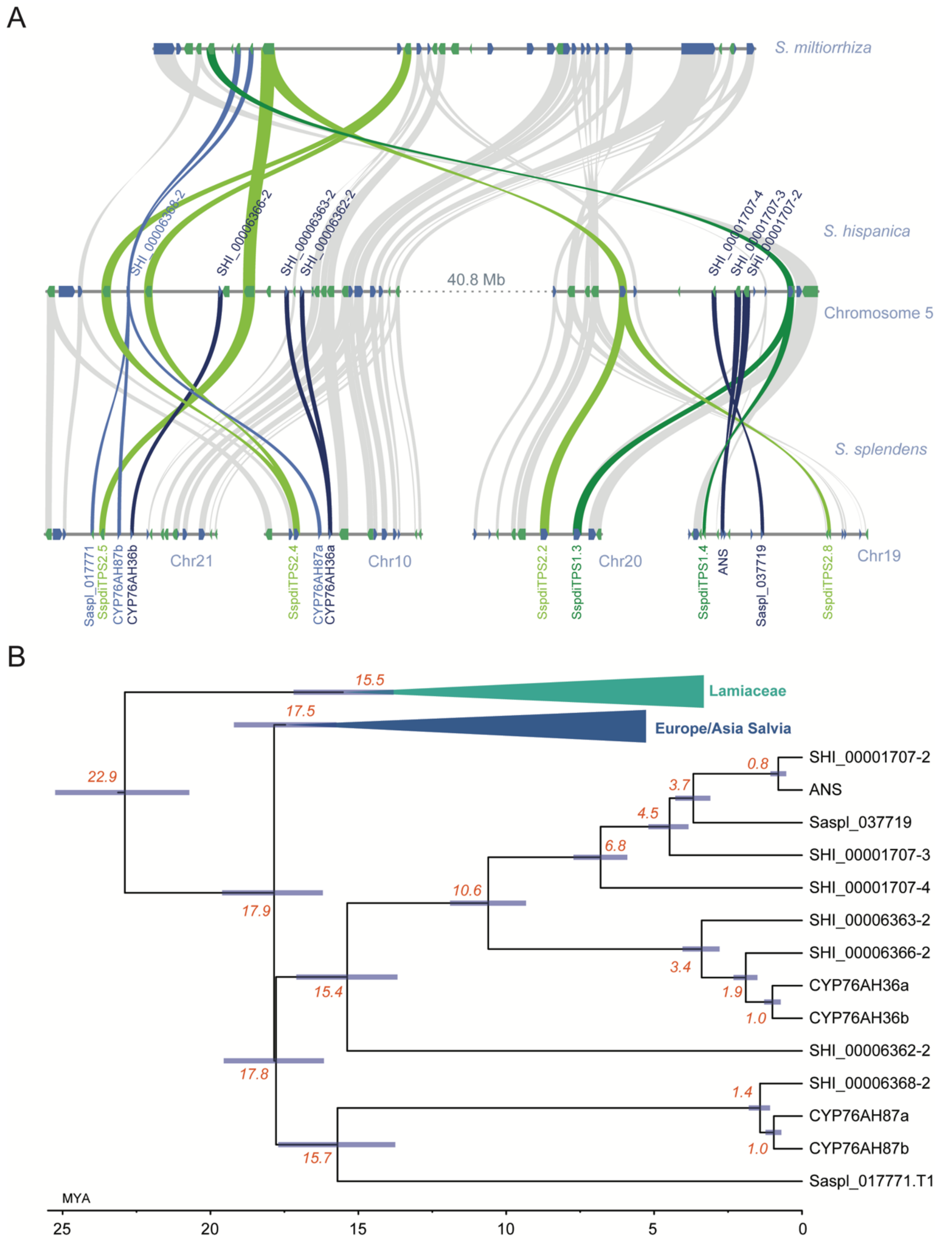


**Fig. S13. Synteny and phylogeny of CYP76AH. A**, Genomic region and syntenic analysis among the neotropical sages *S. splendens*, *S. hispanica* and Asian sage *S. miltiorrhiza*. Dark green and light green ribbons represent class **I** and class **II** diTPS genes, respectively. Light blue ribbons indicate genes linked to CYP76AH87-like genes, while navy blue ribbons highlight CYP76AH36-like genes. **B**, Bayesian phylogeny of syntenicCYP76 genes from panel A. The divergence times are calculated in million years ago **(**MYA).

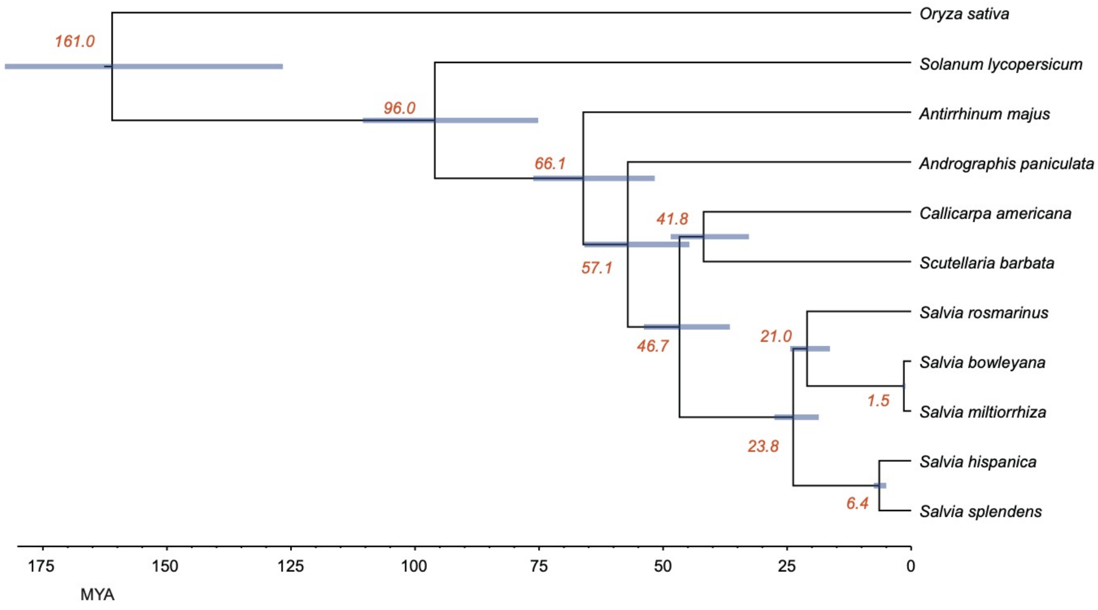


**Fig. S14. Estimated speciation time inferred from single copy ortholog genes across 11 plant genomes.** The 95% confidence interval for the divergence time are marked in blue bars. Estimated divergence times are highlighted in orange fonts. The divergence time of neotropical sages from Eurasia Salvia species is estimated to be around 23.8 million years ago (MYA).

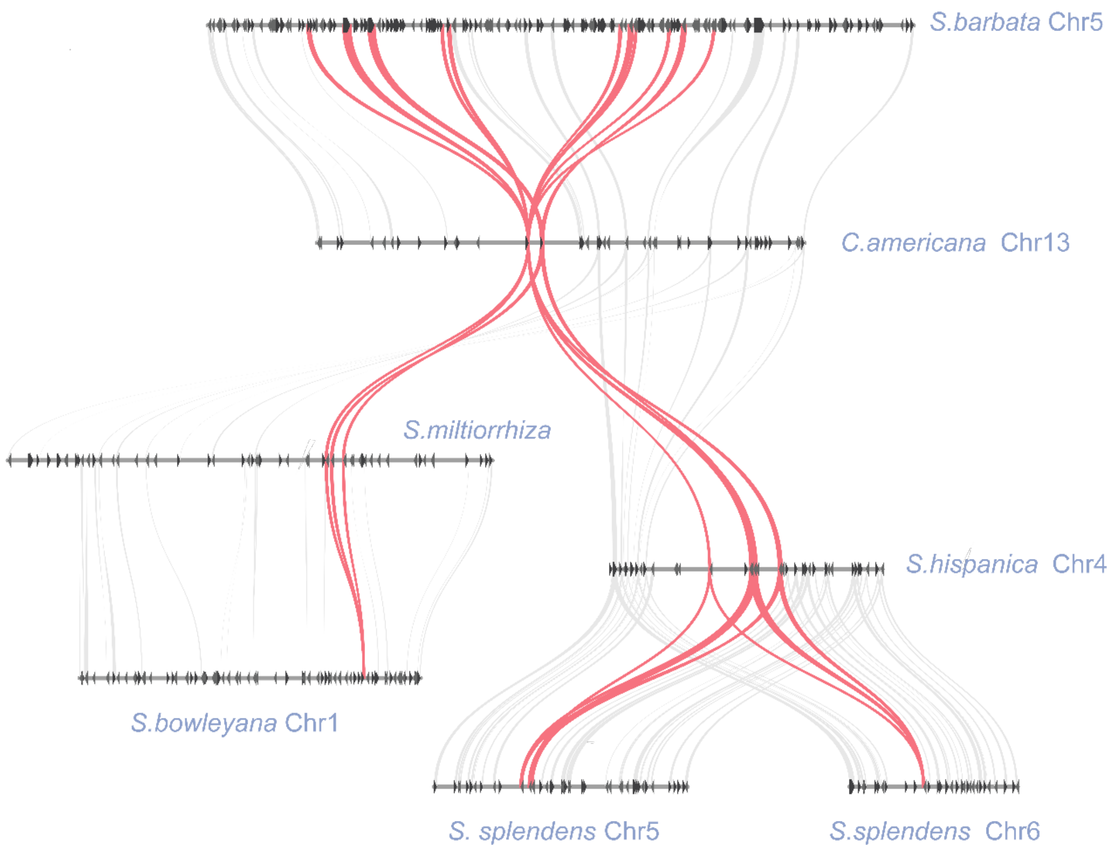


**Fig. S15. Synteny of CYP716B/CYP728D (clan 85) genes.** Syntenic analysis shows that CYP716B genes syntenous to HDAS and HTAS (highlighted by pink ribbon) within family Lamiaceae.

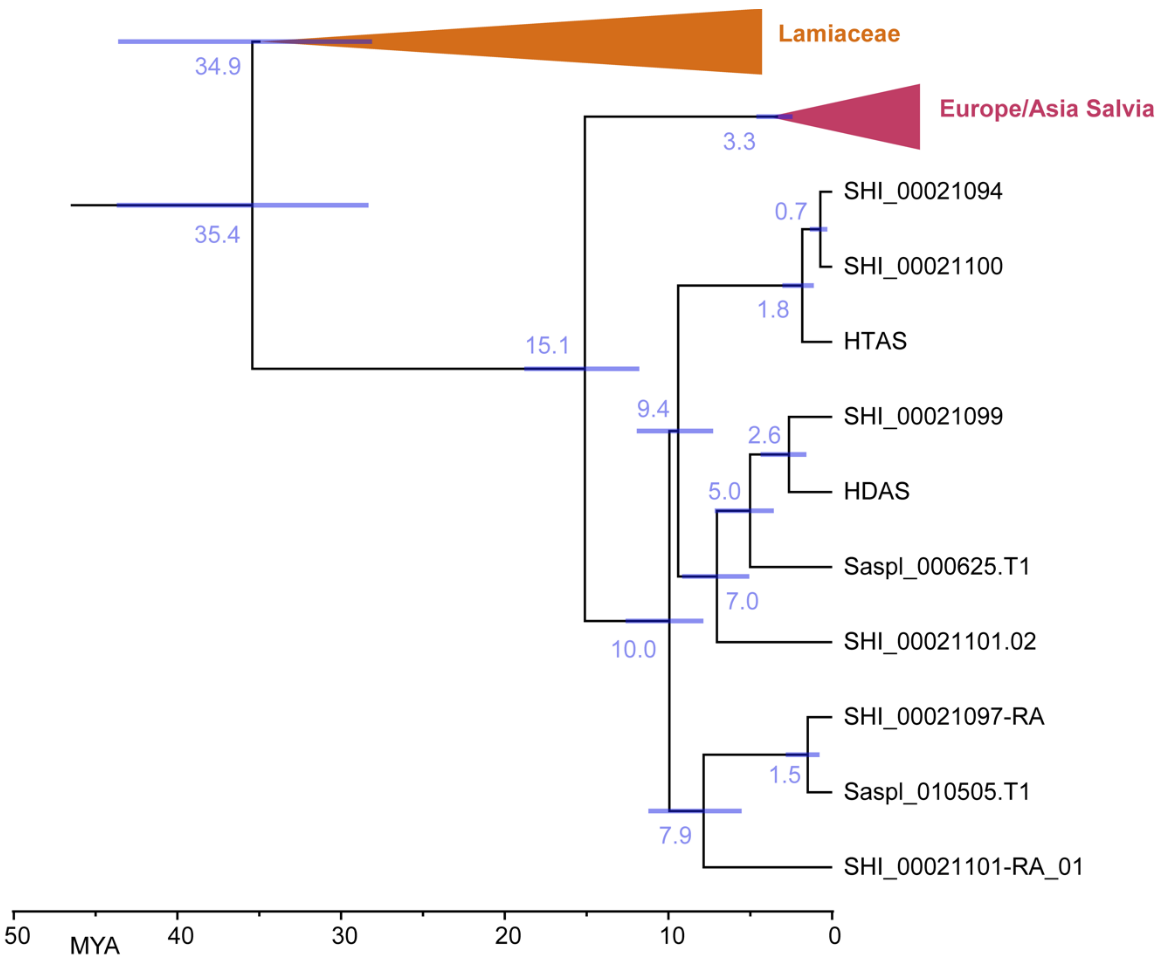


**Fig. S16. Phylogeny of CYP716B/CYP728D (clan 85) in Lamiaceae**. Bayesian phylogeny of syntenous to HDAS and HTAS genes from Fig. S15. The divergence times are calculated in million years ago **(**MYA).

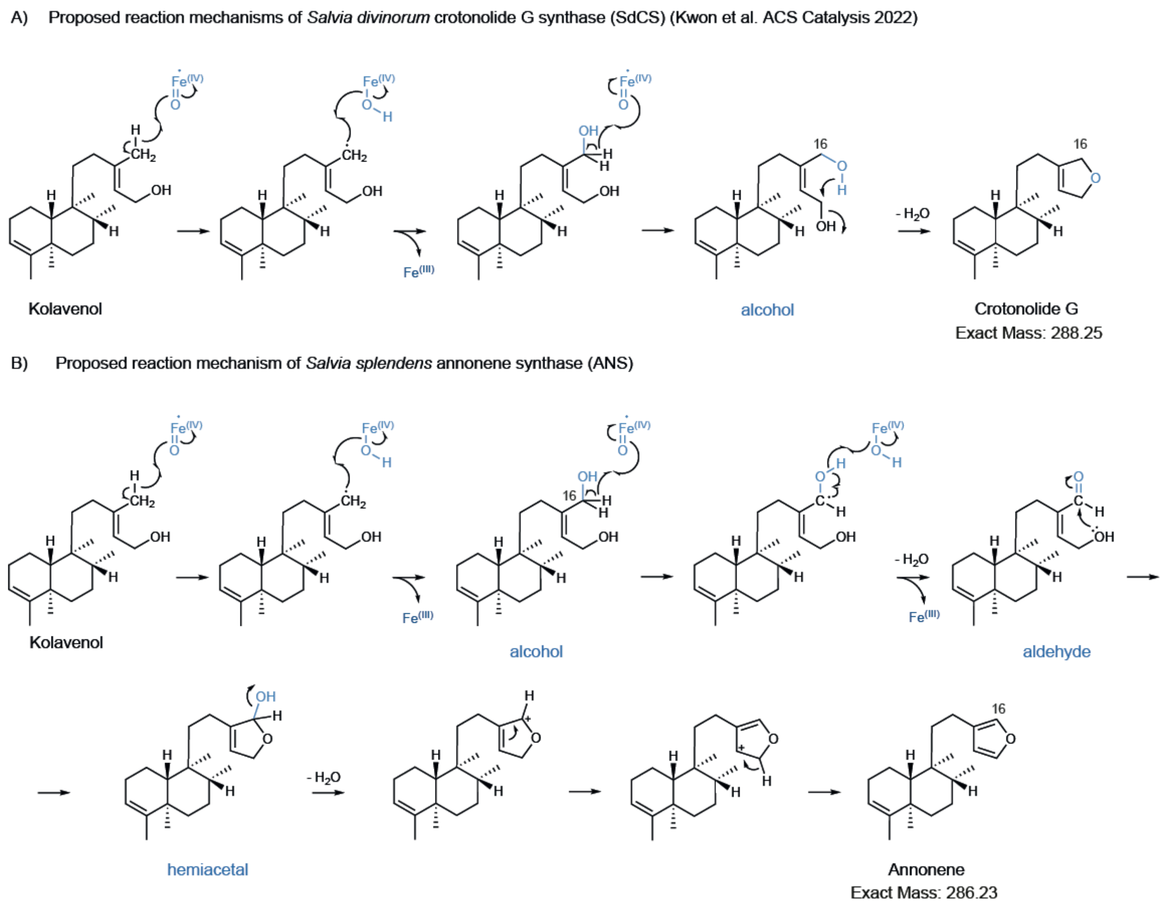


**Fig. S17.** Proposed catalytic mechanism of furan ring on kolavenol (clerodane scaffold) by annonene synthase ANS from *Salvia splendens* in comparison to formation of dihydrofuran ring on kolavenol by the paralogue crotonolide G synthases from *Salvia divinorum (Kwon et al., 2022)*. In the work of Kwon et al. ACS Catalysis 2022 suggest that CYP76AH39 oxidizes the C16 to the corresponding alcohol and subsequently catalyzes the cyclisation to dihydrofuran ring the loss of a water molecule. Annonene synthase subsequently oxidizes C16 to alcohol and then to corresponding aldehyde following the formation of hemiacetal and the final formation of furan ring.

**Table S1. List of genes encoding class I and class II diTPSs and cytochrome P450 enzymes identified from *Salvia splendens* in the present study and our previous work (Li *et al.*, 2023).**

| Protein | Gene ID (scaffold no) | Gene ID (chromosome no) | Activity | NCBI Accession numbers |
|---|---|---|---|---|
| **Class II diTPS** | | | | |
| SspdiTPS2.1 | Saspl_043012 (scaffold82) | SASPL_120943 (chromosome 10) | kolavenyl diphosphate synthase | MT909805 |
| SspdiTPS2.2 | Saspl_048790 (scaffold 372) | SASPL_151044 (chromosome 20) | - | MT909806 |
| SspdiTPS2.3 | Saspl_027494 (scaffold54) | SASPL_135196 (chromosome 13) | *ent*-copalyl diphosphate synthase | MT909807 |
| SspdiTPS2.4 | Saspl_009166 (scaffold 41) | Not annotated (chromosome 10) | copalyl diphosphate synthase | MT909808 |
| SspdiTPS2.5 | Saspl_017770 (scaffold 45) | SASPL_153095 (chromosome 21) | copalyl diphosphate synthase | MT909809 |
| SspdiTPS2.6 | Saspl_009980 (scaffold34) | SASPL_103251 (chromosome 1) | *ent*-copalyl diphosphate synthase | MT909810 |
| SspdiTPS2.7 | Saspl_013955 (scaffold 5) | SASPL_132931 (chromosome 12) | - | MT909811 |
| SspdiTPS2.8 | Saspl_006526 (scaffold 10) | SASPL_149086 (chromosome 19) | - | MT909812 |
| **Class I diTPS** | | | | |
| SspdiTPS1.1 | Saspl_006722 (scaffold 10) | Not annotated (chromosome 17) | ent-kaurene synthase/ kolavenol synthase | MT909800 |
| SspdiTPS1.2 | Saspl_001373 (scaffold 1) | SASPL_146683 (chromosome 18) | ent-kaurene synthase/ kolavenol synthase | MT909801 |
| SspdiTPS1.3 | Saspl_037723 (scaffold 129) | SASPL_149078 (chromosome 19) | miltiradiene synthase/ kolavenol synthase | MT909802 |
| SspdiTPS1.4 | Saspl_048791 (scaffold 372) | SASPL_151043 (chromosome 20) | miltiradiene synthase | MT909803 |
| SspdiTPS1.5 | Saspl_037211 (scaffold 190) | SASPL_127435 (chromosome 10) | kolavenol synthase | MT909804 |
| **P450s** | | | | |
| SspCYP1 | Saspl_009167 (scaffold 41) | SASPL_127726 (chromosome 10) | CYP76AH87a | OP137137 |
| SspCYP2 | Saspl_009168 (scaffold 41) | SASPL_127727 (chromosome 10) | CYP76AH36a ferruginol synthase | OP137138 |
| SspCYP3 | Saspl_017768 (scaffold 45) | SASPL_153093 (chromosome 21) | CYP76AH36b ferruginol synthase | OP137139 |
| SspCYP4 | Saspl_017769 (scaffold 45) | SASPL_153094 (chromosome 21) | CYP76AH87b ferruginol synthase | OP137140 |
| SsANS | Saspl_037721 (scaffold 129) | SASPL_149080 (chromosome 19) | annonene synthase | OR837137 |
| | Saspl_037719 (scaffold129) | SASPL_149084 (chromosome 19) | - | OR837140 |
| SsHDAS | Saspl_010504 (scaffold 24) | SASPL_113922 (chromosome 5) | hardwickiic acid synthase | OR837138 |
| SsHTAS | Saspl_010503 (scaffold 24) | SASPL_113921 (chromosome 5) | hautriwaic acid synthase | OR837139 |
| | Saspl_010505 (scaffold 24) | SASPL_113923 (chromosome 5) | - | OR837141 |

**Table S2. Primers used in this study.**

| Gene name or Gene ID | Primer name | Sequence (5' to 3') |
|---|---|---|
| **Primers used for cloning** | | |
| Saspl_010503 | pESC-Gal1 F | aggagaaaaaaccccggatccgATGGAGTCG ACGATGATTATG |
| | pESC-Gal1 R | caacttctgttccatgtcgacCAACAATTTTTCT GAGGGGTTG |
| Saspl_010503 | pESC-Gal10 F | cactaaagggcggccgcactagtaATGGAGTCG ACGATGATTATG |
| | pESC-Gal10 R | cttgtaatccatcgatactagtgcCAACAATTTTT CTGAGGGGTTG |
| Saspl_010504 | pESC-Gal1 F | aggagaaaaaaccccggatccgATGGAGTCG ACGACGATATT |
| | pESC-Gal1 R | caacttctgttccatgtcgacTCATGCTTTATAA GGTTCTCTGC |
| Saspl_010505 | pESC-Gal1 F | aggagaaaaaaccccggatccgATGATTTTGC TGTGGGTCC |
| | pESC-Gal1 R | caacttctgttccatgtcgacGAATTTGATTGCA TCGTGTAATG |
| Saspl_009167 | pESC-Gal10 F | cactaaagggcggccgcactagtaATGGAGTTG TCCACTGTTGC |
| | pESC-Gal10 R | cttgtaatccatcgatactagtgcTCAATGGCTAA GGGGATAAGC |
| Saspl_009168 and Saspl_017768 | pESC-Gal10 F | cactaaagggcggccgcactagtaATGGACTCC TTCCCTTTCC |
| | pESC-Gal10 R | cttgtaatccatcgatactagtgcTTGCTTAAATG GGATGACC |
| Saspl_037719 | pESC-Gal10 F | cactaaagggcggccgcactagtaATGGATTTC TTCCCTTTCCTG |
| | pESC-Gal10 R | cttgtaatccatcgatactagtgcTGCCTTCTTAT ATGGGATGAC |
| Saspl_037721 | pESC-Gal10 F | cactaaagggcggccgcactagtaatggattcctccttc ccttt |
| | pESC-Gal10 R | cttgtaatccatcgatactagtgctgccttcttgtatgggat ga |
| Saspl_017769 | pESC-Gal10 F | cactaaagggcggccgcactagtaATGGAGTTG TCCACTCTTGC |
| | pESC-Gal10 R | cttgtaatccatcgatactagtgcTCAATGGCTAT GGGGATAAGC |
| SmCYP76AH (SmFS) | pESC-Gal10 F | accctcactaaagggcggccgcaaccATGGATT CTTTTCCTCTCCTC |
| | pESC-Gal10 R | gtcatccttgtaatccatcgatacAGACTTAACTA TTGGGATAATC |
| **Kolavenol production in yeast** | | |
| *ERG20*::F96C | pESC-Gal1 F | aggagaaaaaaccccggatccgATGGCTTCA GAAAAAGAAATTAG |
| | pESC-Gal1 R | caacttctgttccatgtcgacCTATTTGCTTCTC TTGTAAA |
| | F96C mutation F | GCAGGCTTACTGCTTGGTC |
| | F96C mutation R | GACCAAGCAGTAAGCCTGC |

| | | |
|---|---|---|
| SspdiTPS1.5∆50+SspdiTPS2.1 | pESC-Gal10 F | cactaaagggcggccgcactagtaATGTTGGGGGAATTAAAGGACAAGTTGAAGAGGGAAGAA |
| | pESC-Gal10 R | gtcatccttgtaatccatcgatacCTAGACAATTTTTTCAAACA |
| | GGGS linker F | GGTGGTGGTTCTATGAGTATTCAAGCAAACATGTC |
| | GGGS linker R | AGAACCACCACCACTAGGAAGTGTAAGAGGTTCATATA |
| SspdiTPS1.3Δ61+SspdiTPS2.4 | pESC-Gal10 F | cactaaagggcggccgcactagtaATGGAAAACAGTAATTTTCCGGTCACTTTT |
| | pESC-Gal10 R | cttgtaatccatcgatactagtgcCTCGACTGGTTCGAAAAGC |
| | GGGS linker F | GGTGGTGGTTCTATGGCCTCTCTTTCCTCTAC |
| | GGGS linker R | AGAACCACCACCCTTGCCACTCACATTATTAGCT |
| **<u>Heterologous expression of class I diterpene synthases</u>** | | |
| SdKSL1(∆77) pF F | POPIN-F | AAGTTCTGTTTCAGGGCCCGATACCCTCTTATCTGTGTATAATC |
| SdKSL1(∆77) pF R | POPIN-F | ATGGTCTAGAAAGCTTTATTTGCCACTCACATTGTTAGC |
| SdKSL(∆73) pF F | POPIN-F | AAGTTCTGTTTCAGGGCCCGATACCCTCGAGTCTGTGTATAATC |
| SdKSL2(∆73) pF R | POPIN-F | ATGGTCTAGAAAGCTTTAACTCGAAAGTGTAAGAGGCTC |
| **<u>qRT-PCR</u>** | | |
| Saspl_005166 | Ssp qACTIN F | TCAACCCAAAGGCGAATCGT |
| | Ssp qACTIN R | TGACACCATCTCCCGAGTCA |
| Saspl_037721 | Ssp qSsANS F | GCTGGGTTTAATCGAGGGCTA |
| | Ssp qSsANS R | TCGTCTCTGCTCCTCCAACA |
| Saspl_010503 | Ssp qSsHDAS F | GCATTGGCATGACGACCAAG |
| | Ssp qSsHDAS R | ATTGATCGGCACCAGCAACA |
| Saspl_010504 | Ssp qSsHTAS F | GAGATCAGGTTGCACCTCGC |
| | Ssp qSsHTAS R | ACGCACAAAGGTTTCCCTTC |
| Saspl_037211 | Ssp qSsKLS F | CAACGGTTCTTCGTTTGCTCA |
| | Ssp qSsKLS R | AATGGCTTCCTCCTCTGTCAC |
| Saspl_043012 | Ssp qSsKPS F | ATGTAACACTGGTGGCAGAAGG |
| | Ssp qSsKPS R | TCCAATCCGGAGCTCAAATGT |

**Table S3. Vectors used in this study.**

| | Gene vector |
|---|---|
| 1 | pESC-*ERG20*::F96C-SspdiTPS1.5△50+SspdiTPS2.1-His |
| 2 | pESC-*ERG20*::F96C-SspdiTPS1.3△61+SspdiTPS2.4-His |
| 3 | pESC-CrCPR-Leu |
| 4 | pESC-9167-CrCPR-Leu |
| 5 | pESC-9168-CrCPR-Leu |
| 6 | pESC-17768-CrCPR-Leu |
| 7 | pESC-17769-CrCPR-Leu |
| 8 | pESC-37719-CrCPR-Leu |
| 9 | pESC-37721 (SsANS)-CrCPR-Leu |
| 10 | pESC-SmCYP76AH (SmFS)-CrCPR-Leu |
| 11 | pESC-SdCYP76AH39-CrCPR-Leu |
| 12 | pESC-10503 (SsHDAS)-URA |
| 13 | pESC-10504 (SsHTAS)-URA |
| 14 | pESC-10505-URA |
| 15 | pESC-SdCYP728D25-URA |
| 16 | pESC-SdCYP728D26 (SdHDAS)-URA |
| 17 | pESC-SdCYP728D27-URA |

**Dataset S1 (separate file).** Self-organizing map (SOM) and Mutual Ranking (MR) data
